# White Matter Disruption in Pediatric Traumatic Brain Injury: Results from ENIGMA Pediatric msTBI

**DOI:** 10.1101/2020.08.06.237271

**Authors:** Emily L Dennis, Karen Caeyenberghs, Kristen R Hoskinson, Tricia L Merkley, Stacy J Suskauer, Robert F Asarnow, Talin Babikian, Brenda Bartnik-Olson, Kevin Bickart, Erin D Bigler, Linda Ewing-Cobbs, Anthony Figaji, Christopher C Giza, Naomi J Goodrich-Hunsaker, Cooper B Hodges, Elizabeth S Hovenden, Andrei Irimia, Marsh Königs, Harvey S Levin, Hannah M Lindsey, Jeffrey E Max, Mary R Newsome, Alexander Olsen, Nicholas P Ryan, Adam T Schmidt, Matthew S Spruiell, Benjamin SC Wade, Ashley L Ware, Christopher G Watson, Anne L Wheeler, Keith Owen Yeates, Brandon A Zielinski, Peter Kochunov, Neda Jahanshad, Paul M Thompson, David F Tate, Elisabeth A Wilde

**Author notes:** **Please address correspondence to**: Dr. Emily L Dennis, TBI and Concussion Center, Dept of Neurology, University of Utah School of Medicine.

## Abstract

Annually, approximately 3 million children around the world experience traumatic brain injuries (TBIs), of which up to 20% are characterized as moderate to severe (msTBI) and/or have abnormal imaging findings. Affected children are vulnerable to long-term cognitive and behavioral dysfunction, as injury can disrupt or alter ongoing brain maturation. Post-injury outcomes are highly variable, and there is only limited understanding of how inter-individual differences in outcomes arise. Small sample sizes have also complicated efforts to better understand factors influencing the impact of TBI on the developing brain. White matter (WM) disruption is a critical aspect of TBI neuropathology and diffusion MRI (dMRI) is particularly sensitive to microstructural abnormalities. Here we present the results of a coordinated analysis of dMRI data across ten cohorts from three countries. We had three primary aims: (1) to characterize the nature and extent of WM disruption across key post-injury intervals (acute/subacute - within 2 months, post-acute - 2-6 months, chronic - 6+ months); (2) evaluate the impact of age and sex on WM in the context of injury; and (3) to examine associations between WM and neurobehavioral outcomes. Based on data from 507 children and adolescents (244 with complicated mild to severe TBI and 263 control children), we report widespread WM disruption across all post-injury intervals. As expected, injury severity was a significant contributor to the pattern and extent of WM degradation, but explained less variance in dMRI measures with increasing time since injury, supporting other research indicating that other factors contribute increasingly to outcomes over time. The corpus callosum appears to be particularly vulnerable to injury, an effect that persists years post-TBI. We also report sex differences in the effect of TBI on the uncinate fasciculus (UNC), a structure with a key role in emotion regulation. Females with a TBI had significantly lower fractional anisotropy (FA) in the UNC than those with no TBI, and this phenomenon was further associated with more frequent parent-reported behavioral problems as measured by the Child Behavior Checklist (CBCL). These effects were not detected in males. With future harmonization of imaging and neurocognitive data, more complex modeling of factors influencing outcomes will be possible and help to identify clinically-meaningful patient subtypes.

## Introduction

Traumatic brain injury (TBI) can have devastating and long-lasting consequences for brain health and associated functional outcomes, and can be especially disruptive during development in children and adolescents. The plasticity of the human brain during development enables learning and adaptation, but its hidden cost may be increased vulnerability to injury (Anderson *et al.*, 2011; Ismail *et al.*, 2017). Brain injury may derail developmental processes and can drain the neural resources required for typical brain maturation (Hebb, 1949; Pascual-Leone *et al.*, 2005). Children and adolescents also have a higher risk of sustaining a TBI than adults, and TBI is the leading cause of death and disability in youth in the United States (Langlois *et al.*, 2006) and is a significant public health issue worldwide (Dewan *et al.*, 2016). A number of developmental and physiological factors make children particularly vulnerable to poor outcomes (Figaji, 2017). Children have larger head-to-body ratios and less neck strength, which can influence how mechanical injury forces impact the brain. Additionally, incomplete myelination leads to differences in tissue viscosity which can influence response to injury (Guo *et al.*, 2019). TBI can lead to cognitive impairment in all patients, but in children this disruption can become more apparent over time, as children fail to meet the increasing demands of educational activities (Ewing-Cobbs *et al.*, 2004; Anderson *et al.*, 2005; Babikian and Asarnow, 2009; Wells *et al.*, 2009; Ryan *et al.*, 2015).

Advanced magnetic resonance imaging (MRI) techniques have shown great promise in furthering our understanding of the injury and recovery processes that occur during dynamic periods of neurodevelopment (Dennis *et al.*, 2018, Lindsey *et al.*, 2019*b*). Research using diffusion MRI (dMRI) has revealed widespread disruption of brain microstructure in children and adolescents with moderate/severe TBI (msTBI) (Wozniak *et al.*, 2007; Yuan *et al.*, 2007; Levin *et al.*, 2008; Caeyenberghs *et al.*, 2009, 2010, 2012; Oni *et al.*, 2010; Wilde *et al.*, 2010, 2012; Wu *et al.*, 2010; McCauley *et al.*, 2011; Treble *et al.*, 2013, Dennis *et al.*, 2015*a*, *b*, 2017*b*; Johnson *et al.*, 2015; Ewing-Cobbs *et al.*, 2016; Faber *et al.*, 2016; Genc *et al.*, 2017; Königs *et al.*, 2018, Lindsey *et al.*, 2019*a*; Molteni *et al.*, 2019; Watson *et al.*, 2019), which is less likely to be detected by conventional clinical neuroimaging. DMRI can model white matter (WM) tracts and assess tissue structure by mapping the diffusion of water molecules; it is particularly sensitive to traumatic axonal injury, a hallmark of msTBI (Ashwal *et al.*, 2014, Dennis *et al.*, 2017*a*). While this method has shown sensitivity to several forms of TBI-related pathology and often relates to clinical injury and outcome generally, many outstanding questions remain, including the moderating role of key demographic and injury variables, such as age at injury and sex.

Critically, outcomes after TBI are highly variable, and acute clinical measures of injury severity (e.g., Glasgow Coma Scale [GCS], post-traumatic amnesia, loss of consciousness) and conventional clinical neuroimaging only modestly account for inter-individual differences (Anderson *et al.*, 2006; Lajiness-O’Neill *et al.*, 2011, Lindsey *et al.*, 2019*b*; Petranovich *et al.*, 2020). The prefrontal cortex is particularly vulnerable to TBI due to common injury mechanics and to the structure of the skull (Bigler, 2007), potentially contributing to the high prevalence of emotional dysregulation, behavioral change, and executive dysfunction after TBI. Studies have linked atrophy in the corpus callosum (CC) and frontal lobes with several aspects of cognitive dysfunction after TBI (Verger *et al.*, 2001; Slomine *et al.*, 2002; Braga *et al.*, 2007). Around half of children sustaining a msTBI may go on to develop novel psychiatric disorders later in life (Max *et al.*, 2012*a*), although the paucity of longitudinal studies in pediatric msTBI means that more research is needed to better understand factors contributing to psychiatric disorders after TBI, and their prevalence. In fact, there is a significant and specific relationship between novel psychiatric disorders in children with msTBI and WM organization (Max *et al.*, 2012*b*). A more comprehensive understanding of factors that influence outcome post-injury would benefit patients and their families by providing more accurate expectations about recovery and may help to identify additional targets for intervention.

Sex may impact outcome, as some studies have reported longer hospitalizations and greater symptom burden in females (Arambula *et al.*, 2019). Analyses of long-term outcomes have also revealed sex differences in psychosocial dysfunction (Scott *et al.*, 2015). There are significant sex differences in WM development that may influence vulnerability to injury (Ho *et al.*, 2020; Schmied *et al.*, 2020). Pre-clinical evidence also suggests that sex may moderate severity of outcomes (see review by (Gupte *et al.*, 2019)). However, most studies of msTBI in children have not had the sample size necessary to examine this effect (Dennis *et al.*, 2018). Age at injury may also play a role in outcomes as different regions may be more or less vulnerable depending on their stage of maturation (Giza and Prins, 2006), although results from neuroimaging analyses have yielded conflicting results (Ewing-Cobbs *et al.*, 2016, Dennis *et al.*, 2017*b*). Progress to date has been limited by heterogeneity among patients (e.g., severity and nature of injury; age at injury; time since injury; pre- and comorbid factors; access to, quality, and effectiveness of treatment; and psychosocial factors), along with limited sample sizes. This is partially related to the difficulty of recruiting and studying this population and to the challenges inherent in multi-site MRI research (e.g., lack of harmonization across MRI systems and scanners; inherent variation in demographic characteristics between cohorts).

The ENIGMA Diffusion Tensor Imaging (DTI) workflow (Jahanshad *et al.*, 2013) has revealed patterns of altered WM organization across a number of clinical populations (Kochunov *et al.*, 2020), including schizophrenia (Kelly *et al.*, 2018), post-traumatic stress disorder (Dennis *et al.*, 2019), and 22q11.2 Deletion Syndrome (Villalón-Reina *et al.*, 2019). Here we applied these novel analytic methods to pediatric msTBI by analyzing data from over 500 participants across 10 cohorts. We examined alterations in WM organization in three post-injury intervals: acute/sub-acute (within 2 months of injury), post-acute (2-6 months post-injury), and chronic (more than 6 months post-injury). We hypothesized that widespread disruptions in WM organization would be evident in the msTBI group across multiple times post-injury and that key demographic factors such as age and sex would moderate outcome.

## Materials and Methods

### Study Design/Context

The ENIGMA Pediatric msTBI Working Group is a subgroup of the ENIGMA Brain Injury Working Group (Wilde *et al.*, 2020; Dennis *et al.*, 2020), an international collaboration among neuroimaging researchers focused on TBI (Dennis *et al.*, 2020). The strategy behind this collaboration is to leverage the existing framework of the Enhancing NeuroImaging Genetics through Meta-Analysis (ENIGMA) Consortium (Thompson *et al.*, 2020) to answer questions that can only be addressed with larger samples than any single institution or research group can access individually. Through harmonized data processing and meta-analysis, we aim to ensure adequate statistical power to address these questions.

### Study Samples

The ENIGMA Pediatric msTBI dMRI analysis included ten cohorts from seven research sites across three countries, totaling 244 children and adolescents (170 males/74 females, aged 5-20 years) with complicated mild (GCS>13 but abnormal imaging findings), moderate (GCS 9-12), or severe TBI (GCS 3-8) and 263 control children and adolescents (150 males/113 females, aged 5-20 years). The control sample included both healthy controls (HC) and children with orthopedic injuries (OI). Some evidence suggests that these comparison groups differ, so collecting both HC and OI may be the best design when possible (Wilde *et al.*, 2019). Five studies were longitudinal and six were cross-sectional, yielding 646 scans from 507 participants. **Table 1** provides basic demographic and clinical details on the cohorts. Detailed information on the inclusion and exclusion criteria may be found in **Supplementary Table 1**. All participants provided written or verbal informed assent while parents provided written informed consent approved by local institutional review boards. Apart from one cohort, all sites shared raw imaging data with the central site (University of Utah), where they were processed and analyzed. The remaining site processed, quality checked, and analyzed data according to the same set of standardized scripts (accessible on the ENIGMA website: http://enigma.ini.usc.edu/protocols/dti-protocols/).

**Table 1.**
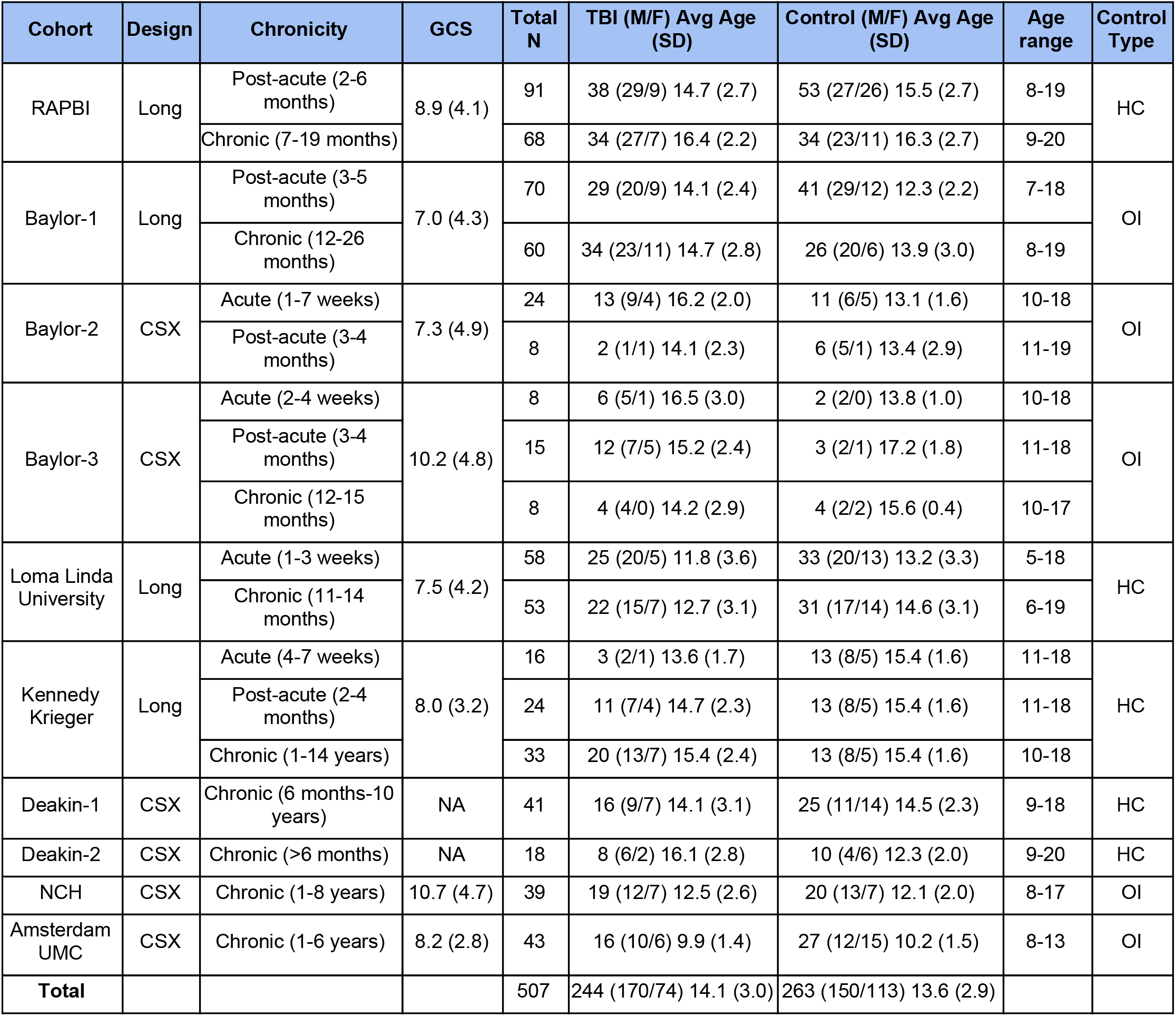
Demographic and clinical details of cohorts. For each cohort, the design (longitudinal - Long, or cross-sectional - CSX), chronicity of injury (acute/sub-acute=less than 2 months post-injury, post-acute=2-6 months post-injury, chronic=more than 6 months post-injury), GCS (average and standard deviation - SD, NA=not available), total N, number of TBI and control participants, number of male and female participants, age range (average and SD), and the type of control group (HC=healthy controls, OI=orthopedically injured) used are listed.

| Cohort | Design | Chronicity | GCS | Total N | TBI (M/F) Avg Age (SD) | Control (M/F) Avg Age (SD) | Age range | Control Type |
| --- | --- | --- | --- | --- | --- | --- | --- | --- |
| RAPBI | Long | Post-acute (2-6 months) | 8.9 (4.1) | 91 | 38 (29/9) 14.7 (2.7) | 53 (27/26) 15.5 (2.7) | 8-19 | HC |
|  |  | Chronic (7-19 months) |  | 68 | 34 (27/7) 16.4 (2.2) | 34 (23/11) 16.3 (2.7) | 9-20 |  |
| Baylor-1 | Long | Post-acute (3-5 months) | 7.0 (4.3) | 70 | 29 (20/9) 14.1 (2.4) | 41 (29/12) 12.3 (2.2) | 7-18 | OI |
|  |  | Chronic (12-26 months) |  | 60 | 34 (23/11) 14.7 (2.8) | 26 (20/6) 13.9 (3.0) | 8-19 |  |
| Baylor-2 | CSX | Acute (1-7 weeks) | 7.3 (4.9) | 24 | 13 (9/4) 16.2 (2.0) | 11 (6/5) 13.1 (1.6) | 10-18 | OI |
|  |  | Post-acute (3-4 months) |  | 8 | 2 (1/1) 14.1 (2.3) | 6 (5/1) 13.4 (2.9) | 11-19 |  |
| Baylor-3 | CSX | Acute (2-4 weeks) | 10.2 (4.8) | 8 | 6 (5/1) 16.5 (3.0) | 2 (2/0) 13.8 (1.0) | 10-18 | OI |
|  |  | Post-acute (3-4 months) |  | 15 | 12 (7/5) 15.2 (2.4) | 3 (2/1) 17.2 (1.8) | 11-18 |  |
|  |  | Chronic (12-15 months) |  | 8 | 4 (4/0) 14.2 (2.9) | 4 (2/2) 15.6 (0.4) | 10-17 |  |
| Loma Linda University | Long | Acute (1-3 weeks) | 7.5 (4.2) | 58 | 25 (20/5) 11.8 (3.6) | 33 (20/13) 13.2 (3.3) | 5-18 | HC |
|  |  | Chronic (11-14 months) |  | 53 | 22 (15/7) 12.7 (3.1) | 31 (17/14) 14.6 (3.1) | 6-19 |  |
| Kennedy Krieger | Long | Acute (4-7 weeks) | 8.0 (3.2) | 16 | 3 (2/1) 13.6 (1.7) | 13 (8/5) 15.4 (1.6) | 11-18 | HC |
|  |  | Post-acute (2-4 months) |  | 24 | 11 (7/4) 14.7 (2.3) | 13 (8/5) 15.4 (1.6) | 11-18 |  |
|  |  | Chronic (1-14 years) |  | 33 | 20 (13/7) 15.4 (2.4) | 13 (8/5) 15.4 (1.6) | 10-18 |  |
| Deakin-1 | CSX | Chronic (6 months-10 years) | NA | 41 | 16 (9/7) 14.1 (3.1) | 25 (11/14) 14.5 (2.3) | 9-18 | HC |
| Deakin-2 | CSX | Chronic (>6 months) | NA | 18 | 8 (6/2) 16.1 (2.8) | 10 (4/6) 12.3 (2.0) | 9-20 | HC |
| NCH | CSX | Chronic (1-8 years) | 10.7 (4.7) | 39 | 19 (12/7) 12.5 (2.6) | 20 (13/7) 12.1 (2.0) | 8-17 | OI |
| Amsterdam UMC | CSX | Chronic (1-6 years) | 8.2 (2.8) | 43 | 16 (10/6) 9.9 (1.4) | 27 (12/15) 10.2 (1.5) | 8-13 | OI |
| **Total** |  |  |  | 507 | 244 (170/74) 14.1 (3.0) | 263 (150/113) 13.6 (2.9) |  |  |

### Image Acquisition and Processing

The acquisition parameters for each cohort are provided in **Supplementary Table 2**. Preprocessing, including eddy current correction, echo-planar imaging-induced distortion correction, and tensor fitting, was performed at the University of Utah. All data were visually quality checked at multiple stages according to the recommended protocols and quality control procedures of the ENIGMA-DTI and NITRC (Neuroimaging Informatics Tools and Resources Clearinghouse) webpages, including careful inspection of registrations. Fractional anisotropy (FA) is a measure of the degree to which water is diffusing preferentially along the direction of axons, and has been interpreted as a proxy for myelin integrity, though it can also be altered by inflammation and axonal packing (Basser *et al.*, 1994). Mean diffusivity (MD) measures the magnitude of diffusion (regardless of direction) in a voxel (averaged across the three eigenvectors), radial diffusivity (RD) is diffusion perpendicular to the largest eigenvalue (typically along the axon), and axial diffusion (AD) is diffusion along the axon. Once tensors were estimated (FA/MD/RD/AD), they were mapped to the ENIGMA DTI template, projected onto the WM skeleton, and averaged within 24 regions of interest (ROIs) from the Johns Hopkins Atlas (JHU), some of which overlap (e.g., genu, body, and splenium of corpus callosum [CC] and total CC, http://enigma.ini.usc.edu/protocols/dti-protocols/). Further details and ROI abbreviations may be found in **Supplementary Note 1**. Across all sites (except the single site that did not share raw imaging data), we extracted motion parameters from the eddy current correction procedure to determine whether motion played a confounding role in our case-control findings. We examined rotation and translation averaged across the X, Y, and Z axes and found greater average rotation (*t*=2.4, *p*=0.018) in the control group. Therefore, we repeated group comparisons while covarying for rotation.

### Statistical Analysis

For each cohort, a linear model was fit using the *lm*, *ppcor,* and *matrixStats* packages in R 3.5.3 (https://www.r-project.org/), with the ROI FA as the response variable and group and covariates as predictors. For cohorts/studies with more than one data collection site, each site’s subjects were analyzed as a separate cohort. As in prior ENIGMA disease working group meta-analyses (Kelly *et al.*, 2018), a random-effects inverse-variance weighted meta-analysis was conducted at a central coordinating site (the University of Southern California Imaging Genetics Center) in R (*metafor* package, version 1.99–118 http://www.metafor-project.org/) to combine individual cohort estimated effect sizes. Cohen’s *d* for the main effect of group and unstandardized *β* coefficients (regression parameters) for continuous predictors were computed with 95% confidence intervals. We used the Cohen’s *d* calculation which accounts for covariates in the fixed effects model, using the following equation:

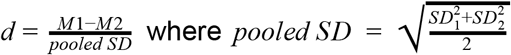

Heterogeneity scores (*I*^2^) for each test were computed, indicating the percent of total variance in effect size explained by heterogeneity across cohorts. As the most commonly-reported dMRI metric, bilaterally-averaged FA was the primary imaging measure, with corresponding MD, RD, and AD examined *post hoc* when FA was significant for an effect of TBI. Lateralized ROIs were examined *post hoc* when a significant effect was found for the bilateral average. The corticospinal tract was not analyzed as its FA measurements have poor reliability, likely due to registration issues (Jahanshad *et al.*, 2013). The average correlation in FA between all pairs of ROIs was *r* =.56. A Bonferroni correction is considered too conservative when there are correlations among the multiple dependent measures being tested (Nyholt, 2004; Li and Ji, 2005). Therefore, we followed recent ENIGMA analyses (Dennis *et al.*, 2019) and calculated the effective number of independent tests based on the observed correlation structure between the regional measures. The equation of Li and Ji (Li and Ji, 2005) yielded Veff = 10, giving a significance threshold of *p*<0.05/10=0.005.

### Code and Data Availability

All analyses were conducted using generalizable scripts available on the ENIGMA GitHub repository: https://github.com/ENIGMA-git/ENIGMA/tree/master/WorkingGroups/EffectSize_and_GLM. Individual ROI-level data were processed using a set of R scripts with regressions customized for the current ENIGMA Pediatric msTBI dMRI analysis workflow, which is available on a set of Google Spreadsheet configuration files by request. Data are available to researchers who join the working group and submit a secondary analysis proposal to the group for approval.

### Non-linear Age Term

We first conducted analyses to examine whether a nonlinear age term should be included in statistical models along with age and sex, as increases and decreases in FA over the lifespan do not follow a linear trend (Kochunov *et al.*, 2010). Age^2^ was significantly associated with FA for a number of ROIs so it was included in all subsequent models.

### Primary Group Comparisons

Data were binned into three post-injury intervals: acute to subacute (MRIs acquired 1 week-2 months after injury), post-acute (2-6 months post-injury), and chronic (6 months-14 years post-injury) (Dennis *et al.*, 2017*a*). Within each of these time periods, we compared groups of patients with TBI and controls. Sites with fewer than 5 participants in any cell were not included in meta-analyses. Six cohorts collected data on HC, while five studies recruited children with OIs (matched for time since injury to the TBI group) as controls. To examine the impact of control group, all group comparisons were repeated separately for those cohorts that recruited HC or OI comparison groups.

### Interactions

We examined potential interactions between group and age or sex, within the three post-injury windows.

### Injury Variables

Within the msTBI group, we examined linear relationships using regression analyses between dMRI measures and three injury variables: age-at-injury (controlling for age-at-scan), GCS, and time since injury (TSI).

### Neurobehavioral Measures

Six of the cohorts collected the parent version of the Behavior Rating Inventory of Executive Function (BRIEF) (Gioia *et al.*, 2000), although two cohorts had too few participants (<5) with both BRIEF and high-quality dMRI to be included in analyses. Among these, we conducted linear regressions on the normative T scores from two summary indices (Behavioral Regulation Index [BRI] and Metacognition Index [MI]) and the Global Executive Composite (GEC) within the TBI group. The BRI assesses behavior that is considered to be related to inhibition, shifting, and emotional control, while the MI assesses behavior considered to be related to the ability to plan, initiate, and monitor activity and performance along with working memory. GEC is a measure of behavior considered to be related to overall executive functioning. There was insufficient data to examine associations between WM organization and BRIEF scores in the acute phase sample. In the post-acute phase sample, 56 participants in the TBI group had BRIEF data. The average **μ**, standard deviation **σ**, and range of the T scores were: for BRI - **μ**=51.9, **σ**=12.0, range=37-79; for MI - **μ**=53.3, **σ**=11.4, range=36-78; for GEC - **μ**=52.8, **σ**=11.8, range=36-74. In the chronic phase, 86 participants in the TBI group had BRIEF data. The average, standard deviation, and range of the T scores were: for GEC - **μ**=51.2, **σ**=10.5, range=32-76; for BRI - **μ**=50.5, **σ**=10.7, range=36-77; for MI - **μ**=51.5, **σ**=10.5, range=30-75. Outliers, defined as being more than 3 SDs away from the age-adjusted population mean, were removed (any T score <21 or >79).

Researchers studying four of the cohorts collected the Child Behavior Checklist (CBCL), a parent report of emotional and behavioral functioning (Achenbach, 1994), although one cohort had too few participants with CBCL and dMRI of acceptable quality to be included in analyses. Among these three cohorts, we conducted linear regressions assessing associations with FA on the T scores from three summary indices - Internalizing Problems, Externalizing Problems, and Total Problems. Internalizing Problems encompasses depressive and anxious symptoms along with somatic complaints, while Externalizing Problems covers aggressive behavior and rule-breaking. Total Problems includes both of these, along with social problems, attention problems, and thought problems such as obsessions and compulsions. These were assessed in the chronic phase, as not enough cohorts collected these measures in other phases. Outliers were removed (any T score <21 or >79). There were 69 participants in the TBI group with CBCL data. The average, standard deviation, and range of scores were: Internalizing Problems - **μ**=51.2, **σ**=12.1, range=33-79; Externalizing Problems - **μ**=48.3, **σ**=11.0, range=33-76; Total Problems - **μ**=49.9, **σ**=12.2, range=24-76.

## Results

### Primary Group Comparison

In the acute/subacute phase (38 TBI/44 control participants), post-acute phase (78 TBI/107 control participants), and chronic phase (160 TBI/190 control participants), we found significantly lower FA in the TBI group across a large number of ROIs, particularly central WM tracts and regions (**Table 2**). Effect sizes across ROIs for each time point are shown in **Figure 1** and **Supplementary Videos 1-3**. Forest plots for the sites contributing to the group comparisons are shown in **Figures 2**-**4**. Follow-up analyses including average rotation as a covariate yielded results consistent with our main analyses (for details, see **Supplementary Note 1** and **Supplementary Figure 1**). *Post hoc* analyses of other diffusion metrics revealed higher RD and MD post-acutely and chronically, with higher RD acutely as well. Acutely, AD was significantly lower across ROIs. Post-acutely and chronically, AD was lower in segments of the CC and higher in other ROIs (**Supplementary Figures 2-4**). Generally, significant results for bilateral ROIs were accompanied by significant results in the lateralized ROIs as well.

**Table 2.**
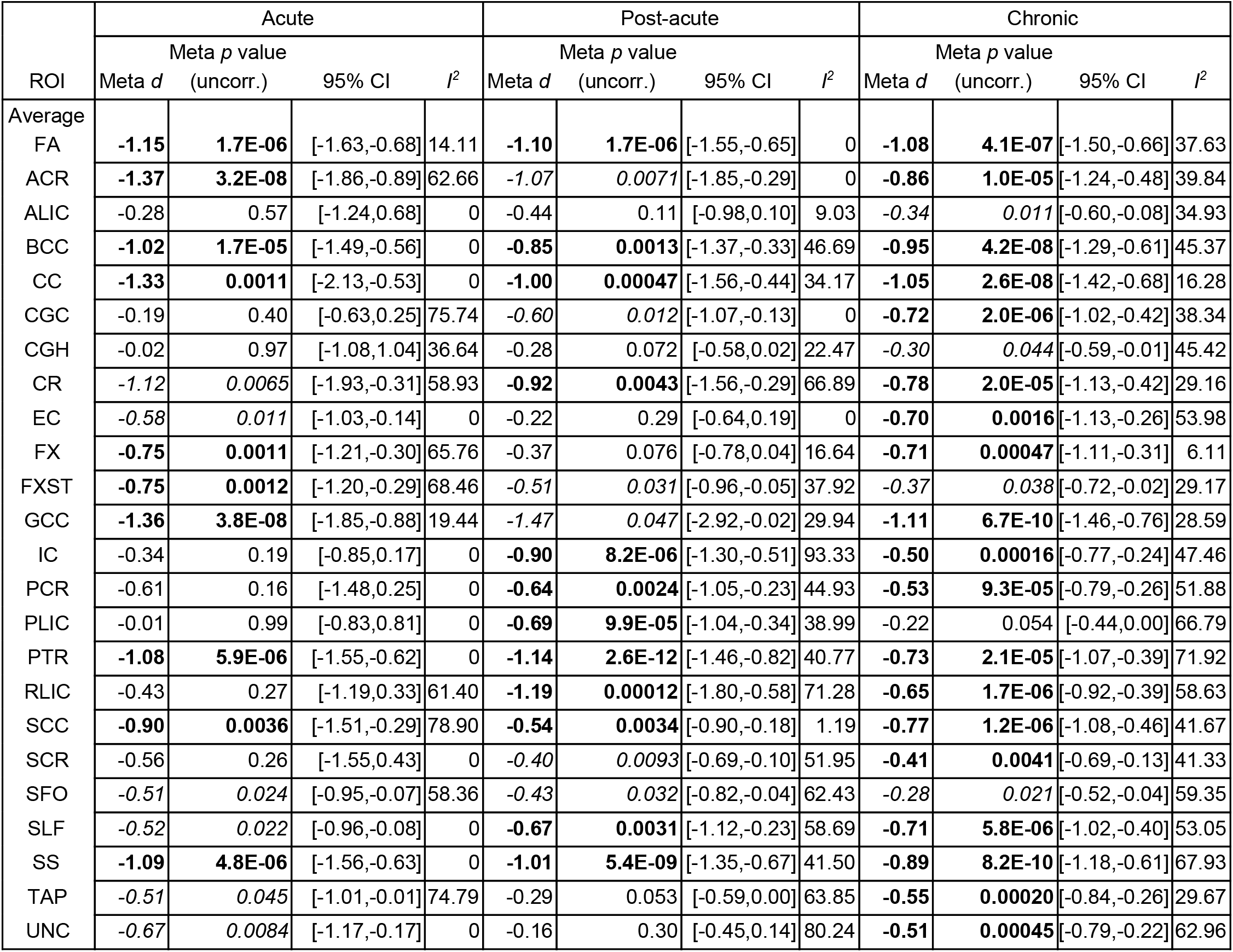
Group differences in FA in the acute/sub-acute, post-acute, and chronic phases. Cohen’s *d* values, uncorrected *p* values, the 95% confidence interval for the *d* statistic, and the *I*^2^ (heterogeneity) are shown for the group comparisons. Bolded results are significant when corrected for multiple comparisons, *italicized* results are marginally significant (based on the Li and Ji adjusted Bonferroni correction, 0.05>*p*>0.005).

**Figure 1.**
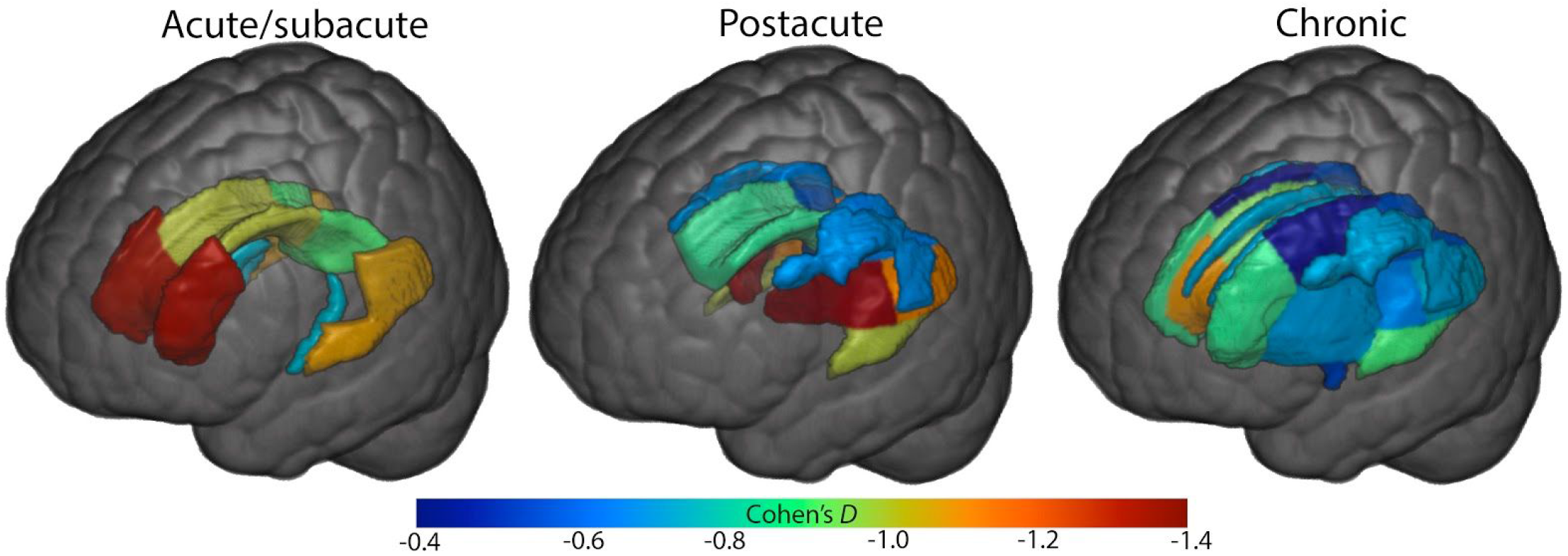
Group differences in FA in the acute/sub-acute, post-acute, and chronic phases. Effect sizes are shown for significant results from the primary group comparison, covarying for sex, age, and age^2^. Cohen’s *D* statistics for midline and bilateral ROIs are displayed according to the color bar below. As TBI was coded as “1” and controls as “0”, negative effect sizes indicate lower FA in the TBI group. Only regions surviving correction for multiple comparisons are shown (*p*<0.005), statistical details for all ROIs are shown in **Table 2**.

**Figure 2.**
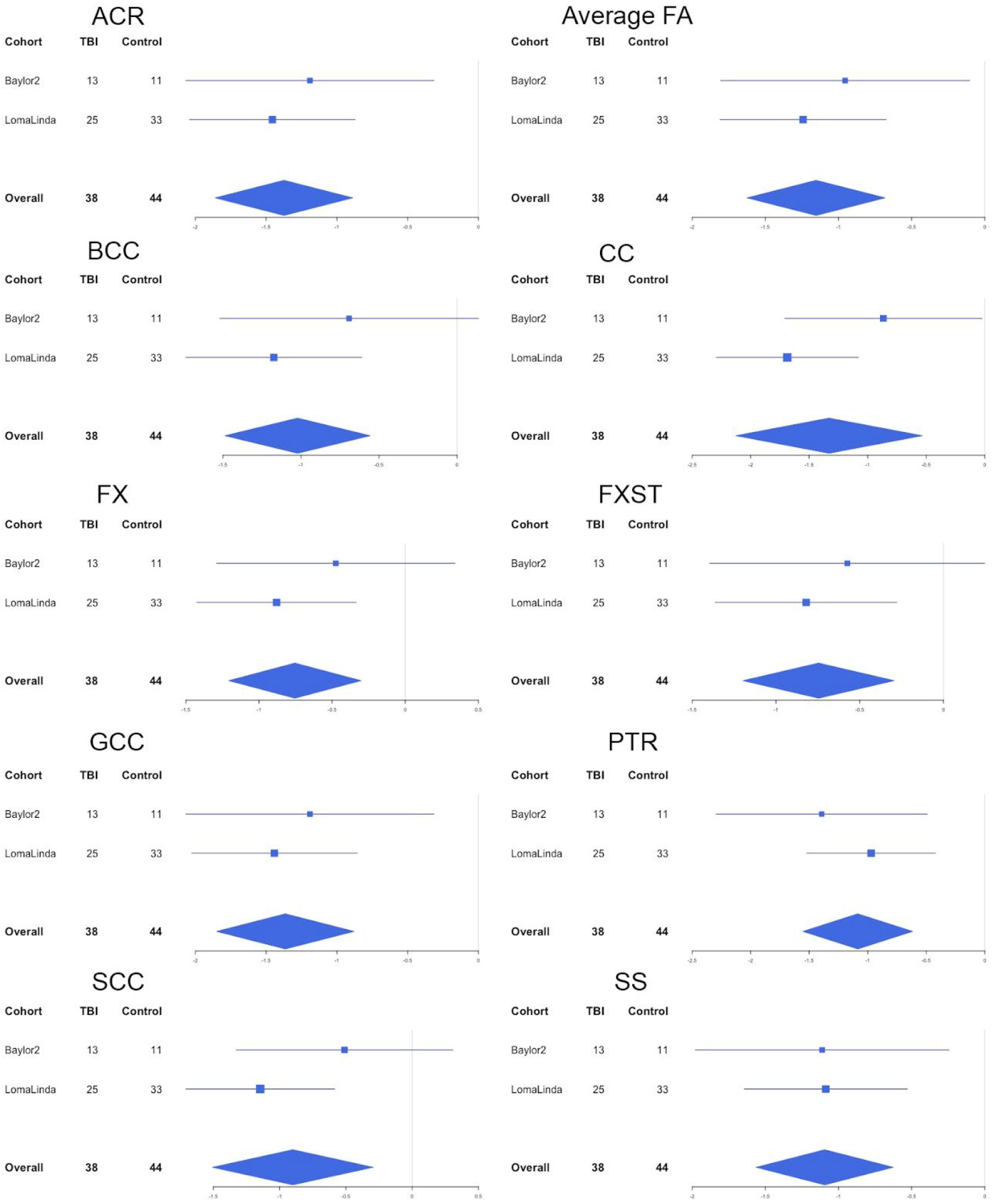
Site effects for the ROIs showing significant group differences in the acute/sub-acute phase. Forest plot shows the effect sizes (Cohen’s D) for the 2 cohorts/sites, scaled by sample size, with bars for 95% CI. The effect size and 95% CI of the meta-analysis is shown at the bottom of the figure.

**Figure 3.**
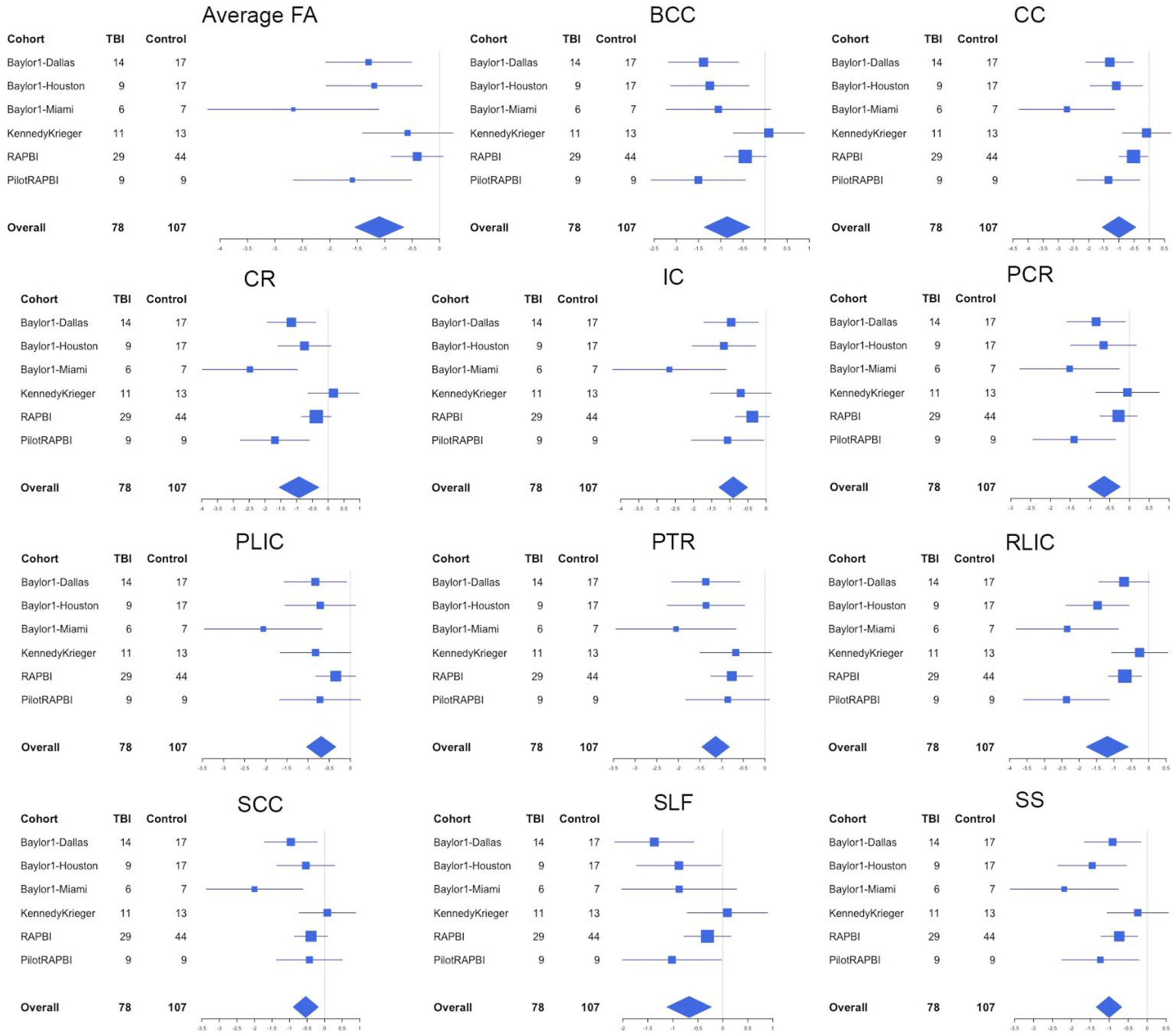
Site effects for the ROIs showing significant group differences in the post-acute phase. Forest plot shows the effect sizes (Cohen’s D) for each of the 6 cohorts/sites, scaled by sample size, with bars for 95% CI. The effect size and 95% CI of the meta-analysis is shown at the bottom of the figure.

**Figure 4.**
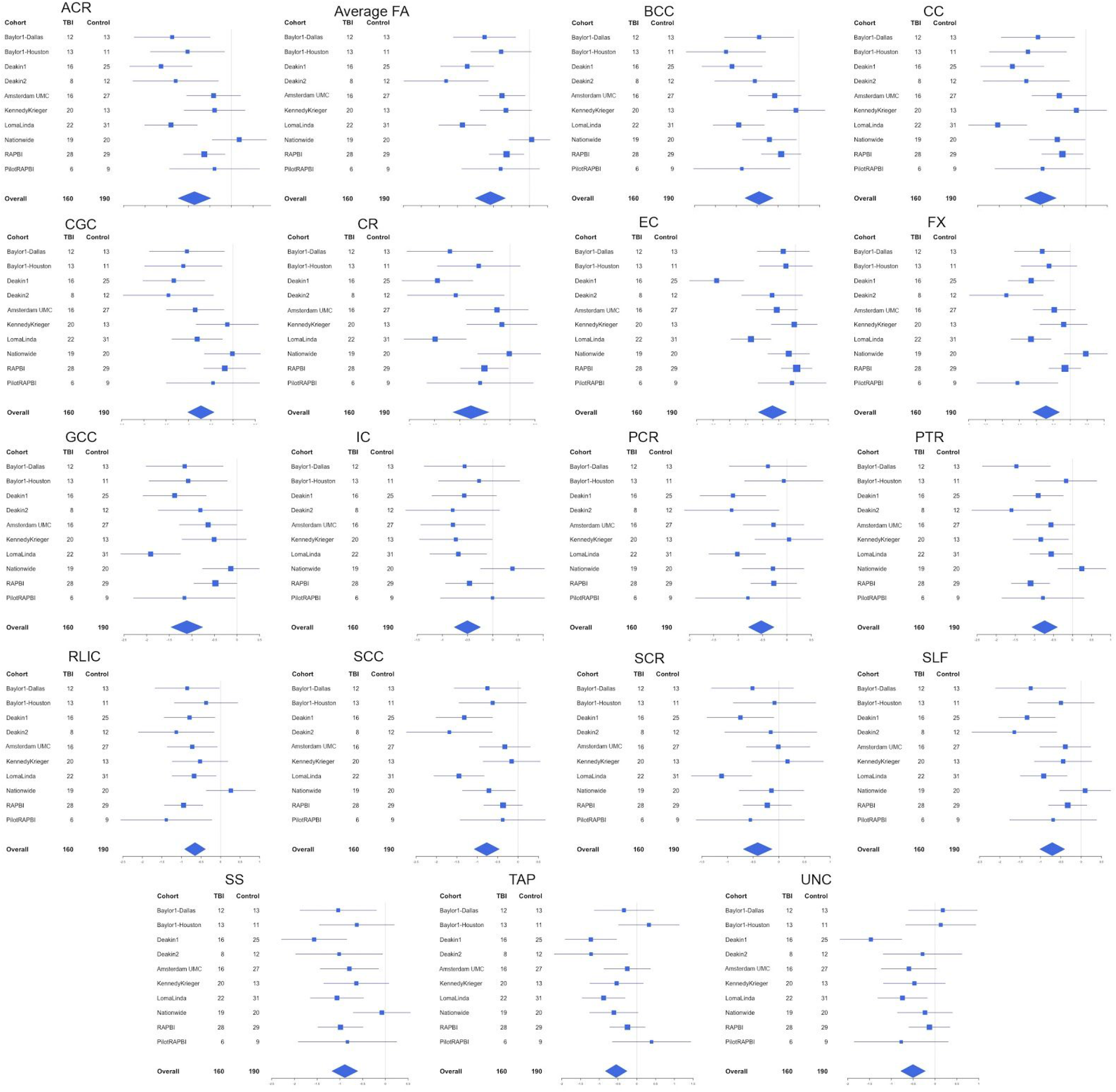
Site effects for the ROIs showing significant group differences in the chronic phase. Forest plot shows the effect sizes (Cohen’s D) for each of the 10 cohorts/sites, scaled by sample size, with bars for 95% CI. The effect size and 95% CI of the meta-analysis is shown at the bottom of the figure.

#### Interactions

A significant group-by-sex interaction was found in the post-acute phase for FA in the UNC (uncinate fasciculus). Further analyses detected no effect of group in males while females with TBI had lower FA than control females (**Figure 5**). In the chronic phase, there were only borderline interaction effects with age or sex (0.005<*p*<0.05, **Supplementary Figure 5**).

**Figure 5.**
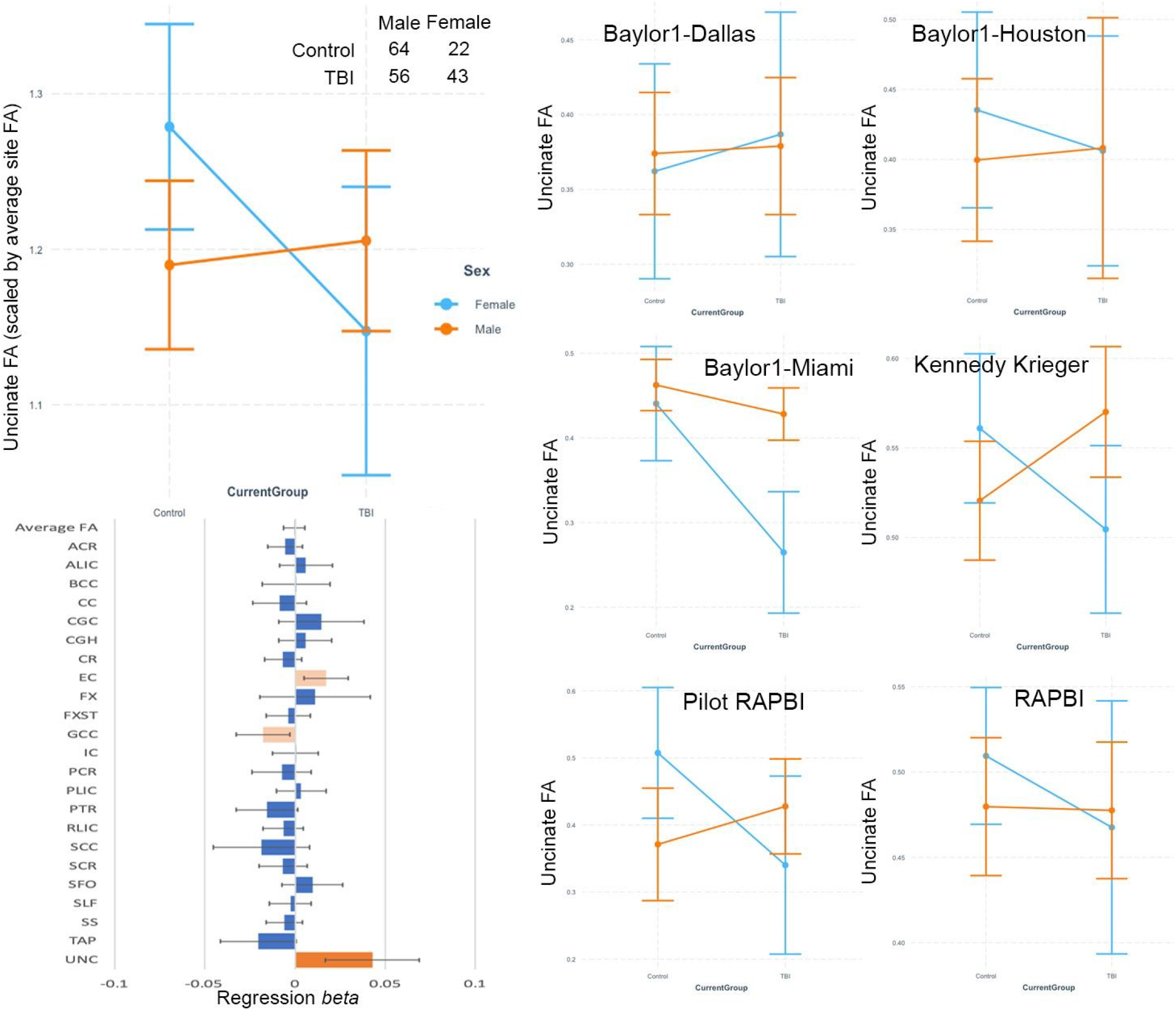
Group-by-sex interactions. Results are shown for the post-acute phase. Shown are unstandardized regression betas for 23 ROIs and average FA (left bottom). Dark orange bars indicate significance (*p*<0.005), light orange bars indicate effects that did not withstand multiple comparisons correction (0.05>*p*>0.005), and blue are not significant (*p*>0.05). Error bars are 95% CI. A plot probing the significant interaction effect in the uncinate is shown in the right panel.

#### Control populations

When conducting separate meta-analyses across sites that recruited HC vs. OI controls, results were generally consistent with the main analyses, although differences were not quite as extensive in the chronic phase for the OI comparison (**Supplementary Figure 6**). There was not a large enough sample (<5 per cell) in the acute phase to examine TBI vs. OI controls.

### Injury Variables

Within the TBI group, significant associations were found with age-at-injury in the post-acute phase in the PTR and SLF (***β***=0.20, *p*=0.00023; ***β***=0.18, *p*=1.3×10^−5^, respectively), with higher FA in patients who were older at the time of injury (**Supplementary Figure 7**). Also post-acutely, significant associations were seen between TSI and the FAs of the BCC and GCC (***β***=−0.0075, *p*=0.00010; ***β***=−0.0049, *p*=0.0041, respectively) with lower FA in patients further from injury (**Supplementary Figure 7**). We found significant associations with GCS within the TBI group at all time points (**Supplementary Figure 7**); in all cases higher GCS (i.e., less severe injury) was associated with higher FA. Acutely, an association was found between GCS and average FA, along with FA of the ALIC, several *corona radiata* (CR) segments, FX, PTR, SCC, SS, and TAP. Post-acutely, GCS was associated with average FA, along with FA of the CR segments, BCC, and SLF. Chronically, GCS was associated with FA of the FX and SS (***β***=0.010, *p*=7.0×10^−6^; ***β***=0.0033, *p*=5.8×10^−5^, respectively).

### Neurobehavioral Function

In the post-acute and chronic phases, respectively, 56 and 86 participants in the TBI group had BRIEF scores. In the post-acute phase, a significant association was found between BRI and average skeleton FA (***β***=−0.00060, *p*=0.0028, **Supplementary Figure 8**), where higher FA was associated with better behavioral regulation. No associations survived correction for multiple comparisons in the chronic phase. For MI, in the chronic phase we found a significant negative association with the FA of the UNC (***β***=−0.0028, *p*=8.6×10^−5^, **Supplementary Figure 8**). No associations survived correction for multiple comparisons in the post-acute phase. For GEC, a number of associations were found in both the post-acute and chronic phase, although none survived correction for multiple comparisons at either time point (**Supplementary Figure 8**).

There were 69 participants across three sites in the TBI group with CBCL scores. Across the TBI group, no significant associations were found between FA and CBCL Internalizing, Externalizing, or Total Problems scores. Given the significant group-by-sex interaction with UNC FA (a key structure for emotion regulation), we also examined the CBCL scores in the female participants in the TBI group only. Across the three sites, 21 female participants in the TBI group had CBCL scores. We found a significant negative association between total problems and FA in the UNC and SS (***β***=−0.0027, *p*=0.0017 and ***β***=−0.0014, *p*=0.0024, respectively, **Figure 6**) with lower FA in patients whose parents reported more problems.

**Figure 6.**
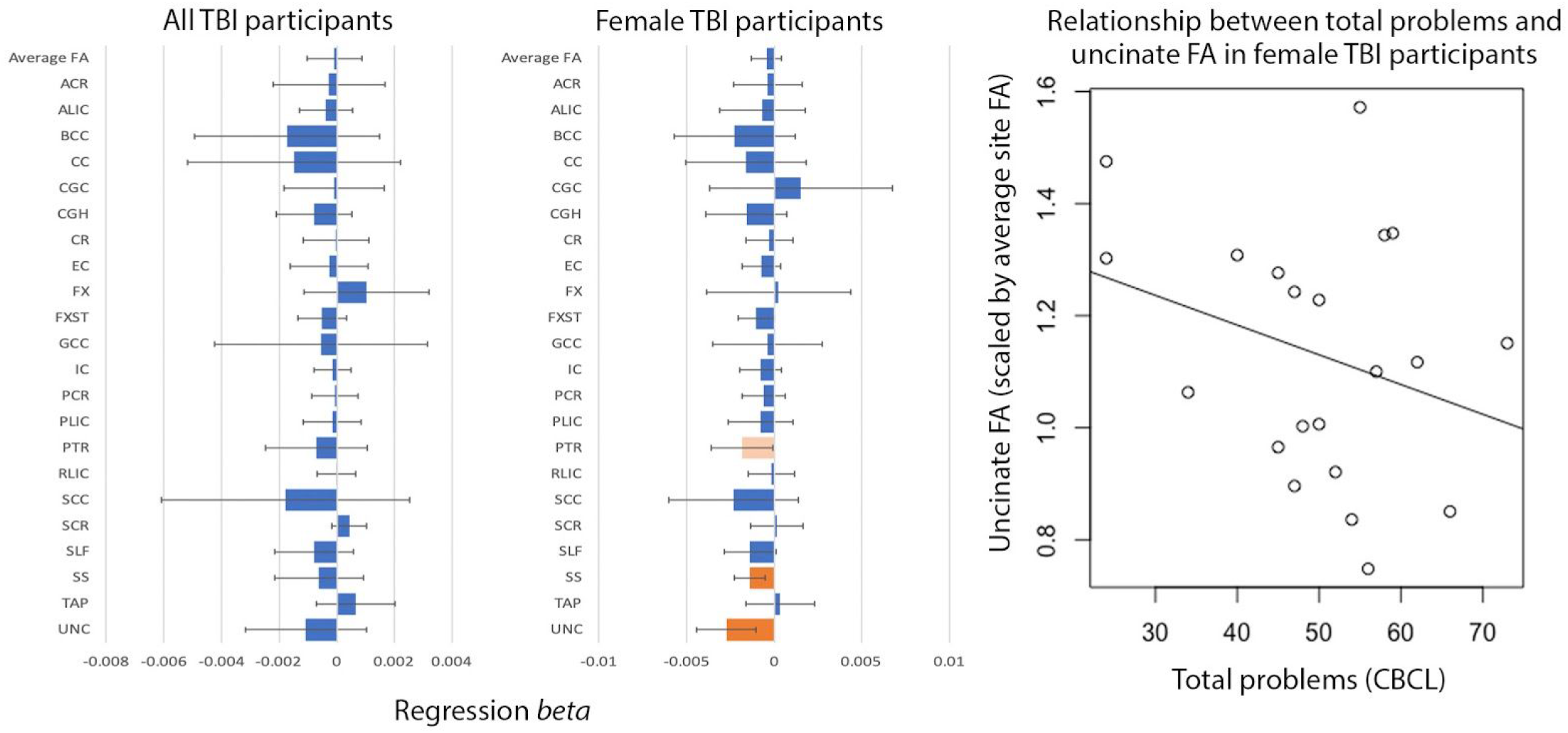
Associations with CBCL Total Problems Score. Linear associations with CBCL Total Problems score in the full TBI group (left) and in the female TBI subset (middle). Shown are unstandardized regression ***β***s for 23 ROIs and average FA. ROI abbreviations are explained in Supplementary Note 1. Dark orange bars indicate significance (*p*<0.005), light orange bars indicate effects that did not withstand multiple comparisons correction (0.05>*p*>0.005), and blue are not significant (*p*>0.05). Error bars are 95% CI. A plot probing the association between total problems and uncinate FA in the female TBI group is shown in the right panel.

## Discussion

Here we present the largest-ever study using dMRI to examine altered WM microstructural organization in pediatric patients with msTBI. In a sample of over 500 children and adolescents from ten cohorts across three countries, we report widespread disruption of WM microstructural organization along all post-injury time windows. We found that females may have a particular vulnerability to WM disruption, especially in the UNC, a fronto-limbic tract, which may underlie a heightened risk of behavioral or emotional problems post-injury. More severe injury is associated with more severe WM disruption, but this effect may lessen with increasing time since injury, evidenced by smaller effect sizes; this may also suggest an increasing influence of other moderating factors over time. Nevertheless, our results indicate that disruption of WM, particularly in callosal fibers, can persist for years post-injury.

In group comparisons, central WM ROIs (CC, CR, internal capsule) exhibited the most extensive disruptions, although, by the chronic phase, nearly every ROI shows significant group differences. This could be for a variety of reasons associated with either pathology and methodology. The CC, in particular, may be most vulnerable to injury as the *falx cerebri* exacerbates lateral forces during an impact (Hernandez *et al.*, 2019). Methodologically, modeling crossing fibers is a known challenge in dMRI that can impact calculations in certain areas like the CR and may mean that alterations in FA are more consistently detected in regions with few mixed fiber populations like the CC. Lower FA, paired with higher MD and RD, could indicate demyelination but could also reflect axonal degeneration, inflammation, or changes in axonal density (Song *et al.*, 2002, Dennis *et al.*, 2017*a*). In the acute/subacute phase, we report lower AD, perhaps reflecting axonal disruption shortly after injury. In the post-acute and chronic phases, however, the directions of AD effects were mixed, with higher AD in the CR and lower AD in the CC. Higher AD could reflect recovery, but it could also result from selective degeneration of neuronal populations. Lower AD in the CC, where the fibers are more unidirectional, suggests axonal degeneration. If callosal projections are interrupted, this could lead to higher AD values in areas where they would have otherwise crossed other fiber bundles, such as the CR. Higher-resolution multi-shell dMRI, which can be used to model intra- and extracellular diffusion, could reveal if neurite density is lower and if there are more unidirectional axonal bundles in the CR further from injury. This would be expected in the presence of selective degeneration of callosal fibers.

In this study, we found a group-by-sex interaction for uncinate fasciculus (UNC) FA. Females with msTBI had lower UNC FA compared to controls, whereas the effect of TBI was not significant in males. The UNC connects the ventral prefrontal cortex and the amygdala and is a key structure for emotion regulation. In prior studies, lower FA in the UNC after TBI was associated with reduced emotional control and increased vulnerability to novel psychiatric disorder, primarily the clinically-significant emotional dysregulation syndrome of personality change due to TBI (Johnson *et al.*, 2011, Max *et al.*, 2012*b*). While data on the long-term outcome after pediatric TBI is relatively sparse, one study reported greater prevalence of internalizing disorders in females compared to males (Scott *et al.*, 2015), although this disparity is also present outside of TBI and may be related to social norms and to sex differences in reporting behaviors (Nolen-Hoeksema, 1990). We also show a significant association between UNC FA and the Total Problems score from the CBCL in female TBI patients. This association was not present in the full TBI group, suggesting that the particular vulnerability of the UNC in girls may lead to a greater likelihood of behavioral or emotional problems after injury. This analysis was underpowered, however, because our sample of females with TBI was small and only three sites collected the CBCL. A central future aim of the ENIGMA Pediatric msTBI working group is harmonizing different scales to extract common domain scores across cohorts, to analyze the neural underpinnings of psychiatric symptoms after TBI in a well-powered, principled manner (Dennis *et al.*, 2020).

Premorbid factors that are associated with brain structure may predispose children to injury (e.g., hyperactivity), and for this reason some studies include OI controls instead of HCs. When we conducted separate meta-analyses of cohorts collecting HC vs. OI controls, results were generally consistent. Using the OI group as a comparison to the TBI group revealed more extensive differences in dMRI in the post-acute phase than when using the HC group as a control. The opposite is true for the chronic phase, although statistical power presumably differed in the chronic phase given the differing sample sizes (chronic phase: 73 TBI vs 77 OI, 100 TBI vs. 119 HC). This disparity could also point to long-term impacts of injury that are not restricted to TBI. Hospitalization, psychological trauma from the injury event, and biological responses associated with secondary injury (such as inflammation) could all contribute to alterations in brain structure and function even when the brain itself is not directly injured (McDonald *et al.*, 2016; Sheeler, 2016; Yang *et al.*, 2016; Nicholson *et al.*, 2018; Ewing-Cobbs *et al.*, 2019).

We examined a number of clinical variables within the TBI group, including age-at-injury, GCS, and time since injury (TSI). Older patients may fare better, as we found significant associations with age-at-injury for the SLF and PTR - two regions that are still maturing throughout adolescence (Schmithorst *et al.*, 2002; Barnea-Goraly *et al.*, 2005; Ashtari *et al.*, 2007; Lebel *et al.*, 2012). However, these associations were only present in the post-acute phase, suggesting that in the long-term the effect is minimal, possibly reflecting late catch-up recovery in younger children with TBI in the chronic phase. TSI similarly showed an effect only in the post-acute phase. This is not surprising, however, as FA calculations in the acute phase may be influenced by acute pathologies such as swelling or the breakdown of the blood-brain barrier (Niogi and Mukherjee, 2010). The lack of detectable associations in the chronic phase (range of post-injury time intervals: 0.5-14 years) may be influenced by variability among cohorts, or it could indicate that the impact of TBI on WM organization may stabilize within the first year or so of injury. Longitudinal studies with more than two assessments are critical to answer this important question. GCS was significantly positively associated with FA at all time points, suggesting that more severe injury is associated with poorer WM organization. These effects were less pronounced in the chronic phase, however, suggesting that other factors besides severity may increasingly account for variations in long-term outcomes. Previous research has found that post-injury family environments influence long-term behavioral outcomes, although these factors have not explicitly been examined with regard to WM organization (Yeates *et al.*, 1997, 2004; Schmidt *et al.*, 2010). Again, more longitudinal studies are needed to show this directly and identify moderating factors.

A limitation of our study is the variability among sites, scan parameters, recruitment criteria, and collected measures. This heterogeneity limits our ability to characterize the groups in great detail and limits our power for some analyses even with our large sample size, particularly those involving behavioral measures. For example, the associations we report with CBCL Total Problems in female TBI patients need to be replicated in a larger sample. However, this discovery was only possible with the relatively large sample that we had, and demonstrates the potential of ENIGMA analyses to generate hypotheses that future research can interrogate in greater depth. The ENIGMA Pediatric msTBI group will conduct follow-up analyses once we establish harmonization procedures that enable us to measure behavioral and psychological disruption across measures (e.g., K-SADS-PL (Kaufman *et al.*, 2013) and VABS (Sparrow *et al.*, 2012)). Another limitation is the inability to control for preinjury behavioral problems and psychiatric diagnoses. The broad variability across sites in the timing of assessments may limit results, as the first year post-injury is especially dynamic from a neural reorganization perspective. We attempted to address this by establishing post-injury intervals but biological changes occur along more of a continuum rather than in discrete periods during recovery from injury, and the scale and granularity of this continuum differ across patients. Variability in study parameters is, to some extent, a strength, as it supports the generalizability of our results. Our analysis includes a good portion of the dMRI data that currently exists for pediatric msTBI cohorts, but it is a small field. Future data collection (ideally with a greater degree of harmonization) will be necessary to move the field forward.

## Conclusion

We conducted a harmonized analysis across cohorts to examine WM organization after TBI. In this analysis of 507 children and adolescents, we report widespread disruption in WM organization following complicated mild to severe TBI. These alterations appear to persist and encompass a larger number of WM regions with time post-injury, although future longitudinal analyses are needed to map true changes in the measures examined, and to identify subsets of patients who fare better. Future analyses employing more complex harmonization and machine learning approaches may reveal clinically-significant patient subtypes based on demographic, clinical, and imaging variables.

## Acknowledgments

K99NS096116 (ELD). R01NS05400 (BB). National Health and Medical Research Council Career Development Fellowship (KC). NRF SARChI Chair of Clinical Neurosciences (AF). UCLA Easton Clinic for Brain Health, Stan and Patty Silver, UCLA Brain Injury Research Center, R01HD061504 (CCG). K01HD083459 (KRH). R01NS100973, DoD contract W81XWH-18-1-0413 and a Hanson-Thorell Research Scholarship (AI). R01HD088438 (JEM). K12HD001097, K23HD06161, UL1TR001079, 1S10OD021648 (SJS). U54EB020403, R01MH116147, R56AG058854, P41EB015922, R01MH111671 (PMT). K99MH119314, NARSAD 27786 (BSCW). Ronald and Irene Ward Chair in Pediatric Brain Injury, funded by the Alberta Children’s Hospital Foundation (KOY). We also wish to acknowledge the participation of the children and family members and the efforts of our many colleagues that make this work possible.

