## Supplementary Material for "White Matter Disruption in Pediatric Traumatic Brain Injury: Results from ENIGMA Pediatric msTBI"

**Supplementary Information**
**Supplementary Note 1.** Further details of ENIGMA-DTI protocols.
**Supplementary Table 1.** Clinical details and inclusion and exclusion criteria for each of the cohorts included.
**Supplementary Table 2.** DTI acquisition parameters for each of the cohorts included.
**Supplementary Video 1.** Significant group differences in the acute/subacute phase. ROIs showing significant group differences are overlaid, with effect sizes corresponding to the color bar.
**Supplementary Video 2.** Significant group differences in the postacute phase. ROIs showing significant group differences are overlaid, with effect sizes corresponding to the color bar.
**Supplementary Video 3.** Significant group differences in the chronic phase. ROIs showing significant group differences are overlaid, with effect sizes corresponding to the color bar.
**Supplementary Figure 1.** Group differences covarying for rotation parameters. Effect sizes are shown for the primary group comparison in the acute, post-acute, and chronic phases, covarying for sex, age, $age^2$, and average rotation. Cohen's *D* statistics are shown across all midline and bilateral ROIs, along with average FA, with bars indicating the 95% confidence interval. As TBI was coded as "1" and controls as "0", negative effect sizes indicate lower FA in the TBI group. ROI abbreviations are explained in Supplementary Note 1. Dark orange bars indicate significance ($p<0.005$), light orange bars indicate effects that did not withstand multiple comparisons correction ($0.05>p>0.005$), and blue are not significant ($p>0.05$).
**Supplementary Figure 2.** Group differences in the acute phase for all four diffusion metrics. Effect sizes are shown for FA, MD, RD, and AD, covarying for sex, age, and $age^2$. Cohen's *D* statistics are shown across all midline and bilateral ROIs, along with average FA, with bars indicating the 95% confidence interval. As TBI was coded as "1" and controls as "0", negative effect sizes indicate lower FA in the TBI group. ROI abbreviations are explained in Supplementary Note 1. Dark orange bars indicate significance ($p<0.005$), light orange bars indicate effects that did not withstand multiple comparisons correction ($0.05>p>0.005$), and blue are not significant ($p>0.05$).
**Supplementary Figure 3.** Group differences in the post-acute phase for all four diffusion metrics. Effect sizes are shown for FA, MD, RD, and AD, covarying for sex, age, and $age^2$. Cohen's *D* statistics are shown across all midline and bilateral ROIs, along with average FA, with bars indicating the 95% confidence interval. As TBI was coded as "1" and controls as "0", negative effect sizes indicate lower FA in the TBI group. ROI abbreviations are explained in Supplementary Note 1. Dark orange bars indicate significance ($p<0.005$), light orange bars indicate effects that did not withstand multiple comparisons correction ($0.05>p>0.005$), and blue are not significant ($p>0.05$).
**Supplementary Figure 4.** Group differences in the chronic phase for all four diffusion metrics. Effect sizes are shown for FA, MD, RD, and AD, covarying for sex, age, and $age^2$. Cohen's *D* statistics are shown across all midline and bilateral ROIs, along with average FA, with bars indicating the 95% confidence interval. As TBI was coded as "1" and controls as "0", negative effect sizes indicate lower FA in the TBI group. ROI abbreviations are explained in Supplementary Note 1. Dark orange bars indicate significance ($p<0.005$), light orange bars indicate effects that did not withstand multiple comparisons correction ($0.05>p>0.005$), and blue are not significant ($p>0.05$).
**Supplementary Figure 5.** Group-by-age and group-by-sex interactions for the acute, post-acute, and chronic phases. Unstandardized regression $\beta$s are shown for 23 ROIs and average FA. ROI abbreviations are explained in Supplementary Note 1. Dark orange bars indicate significance ($p<0.005$), light orange bars indicate effects that did not withstand multiple comparisons correction ($0.05>p>0.005$), and blue are not significant ($p>0.05$). Error bars are 95% CI.
**Supplementary Figure 6.** Group comparisons with healthy controls (HC) and orthopedic injury controls (OI) separated. Effect sizes are shown for FA in the post-acute and chronic phases, covarying for sex, age, and $age^2$. Cohen's *D* statistics are shown across all midline and bilateral ROIs, along with average FA, with bars indicating the 95% confidence interval. As TBI was coded as "1" and controls as "0", negative effect sizes indicate lower FA in the TBI group. ROI abbreviations are explained in Supplementary Note 1. Dark orange

bars indicate significance ($p$<0.005), light orange bars indicate effects that did not withstand multiple comparisons correction (0.05>$p$>0.005), and blue are not significant ($p$>0.05).

**Supplementary Figure 7.** Linear associations with age-at-injury, time since injury, and Glasgow Coma Scale in the TBI group in the acute, post-acute, and chronic phases. Shown are unstandardized regression $\boldsymbol{\beta}$s for 23 ROIs and average FA. ROI abbreviations are explained in Supplementary Note 1. Dark orange bars indicate significance ($p$<0.005), light orange bars indicate effects that did not withstand multiple comparisons correction (0.05>$p$>0.005), and blue are not significant ($p$>0.05). Error bars are 95% CI.

**Supplementary Figure 8.** Linear associations with BRIEF scores in the post-acute and chronic phases. Shown are associations with Global Executive Composite (GEC), Behavioral Regulation Index (BRI), and Meta-Cognition Index (MC). Shown are unstandardized regression $\boldsymbol{\beta}$s for 23 ROIs and average FA. ROI abbreviations are explained in Supplementary Note 1. Dark orange bars indicate significance ($p$<0.005), light orange bars indicate effects that did not withstand multiple comparisons correction (0.05>$p$>0.005), and blue are not significant ($p$>0.05). Error bars are 95% CI.

**Supplementary Note 1. Further details of ENIGMA-DTI protocols.**

Details on scanner and acquisition parameters are provided in **Supplementary Table 2**. Preprocessing, including eddy current correction, EPI induced distortion correction, and tensor fitting, were carried out at each site. Image analysis was conducted at each site using tract-based spatial statistics (TBSS) as part of FSL software (Smith *et al.*, 2006). All data were visually quality checked at multiple stages according to the recommended protocols and quality control procedures of the ENIGMA-DTI and NITRC (Neuroimaging Informatics Tools and Resources Clearinghouse) webpages, including careful inspection of registrations. Participants with large lesions that impacted registration quality were removed from analyses. Individual subject FA maps were aligned to the custom ENIGMA-DTI FA template derived from 400 adult participants scanned across four sites designed for optimal multi-site harmonization (Jahanshad *et al.*, 2013). While a pediatric atlas may be more accurate, using the ENIGMA template ensures comparability with other working groups and has been used in other analyses of pediatric and adolescent cohorts (Piras *et al.*, 2019; van Velzen *et al.*, 2019; Villalón-Reina *et al.*, 2019). Creation of a pediatric template is a future goal. FA voxels were then projected onto the ENIGMA-DTI template skeleton. This creates a unique FA skeleton in the same space for each individual in each cohort. To minimize effects of residual registration misalignment, the regions of interest were consistent in size across sites and the skeletonization procedure was performed individually for each site to minimize any site-specific residual misalignment. The same projection used for the FA images also projects the non-FA (mean, axial, and radial) images onto the skeleton. Voxels along the individual skeletons were averaged across white matter ROIs. A total of 25 bilateral ROIs were delineated based on the JHU WM atlas, an established WM parcellation derived using deterministic tractography (Mori *et al.*, 2008). A whole-brain WM skeleton was defined according to the tract-based spatial statistics methodology and ROI-averaged measures of FA, MD, AD and RD were then calculated by averaging each of these voxel measures over all skeleton voxels encapsulated by a particular ROI. This ensured that voxels at the periphery of a fiber bundle, where residual registration misalignment is typically maximal, were excluded from the ROI average. In other words, ROI averaging was performed based on the core of each fiber bundle, as defined by the WM skeleton.

The multi-subject JHU white matter atlas was used to parcellate regions of interest from the ENIGMA template in MNI space, with updated label identification to correct an earlier atlas error (Rohlfing, 2013). A total of eighteen bilateral white matter ROIs were extracted from the skeletonized FA images and averaged (the corticospinal tract was ignored as prior reports have shown it to have poor reliability). The table below lists 24 ROIs (some partially overlapping) that were extracted from the skeletonized images, including 5 midsagittal regions (no lateralized components), and 18 lateralized regions (left and right were averaged to obtain bilateral FA). The overall average FA values were calculated by averaging values for the entire white matter skeleton.

ENIGMA-DTI QA/QC protocol consists of visual inspection of the images before and after registration to the ENIGMA template, as well as calculating the average skeleton projection distance. The distance of voxel projection to the ENIGMA skeleton can assess the registration quality between individual images and ENIGMA-DTI template. Higher projection distance may indicate problems with aligning the individual brain to the template. After ROI extraction, histograms of FA and diffusivity measures are computed for each ROI.

| **Abbreviation** | **Full tract name** | **Abbreviation** | **Full tract name** |
|---|---|---|---|
| Average FA | Full skeleton average FA | IC (L+R) | Internal capsule |
| ACR (L+R) | Anterior *corona radiata* | PCR (L+R) | Posterior *corona radiata* |
| ALIC (L+R) | Anterior limb of internal capsule | PLIC (L+R) | Posterior limb of internal capsule |
| BCC | Body of *corpus callosum* | PTR (L+R) | Posterior thalamic radiation |
| CC | *Corpus callosum* | RLIC (L+R) | Retrolenticular part of internal capsule |
| CGC (L+R) | Cingulum (cingulate gyrus) | SCC | *Splenium* of *corpus callosum* |

| | | | |
|---|---|---|---|
| CGH (L+R) | Cingulum (hippocampal portion) | SCR (L+R) | Superior *corona radiata* |
| CR (L+R) | *Corona radiata* | SFO (L+R) | Superior fronto-occipital fasciculus |
| EC (L+R) | External capsule | SLF (L+R) | Superior longitudinal fasciculus |
| FX | *Fornix* | SS (L+R) | Sagittal *stratum* |
| FXST (L+R) | *Fornix* (cres) / *Stria terminalis* | UNC (L+R) | *Uncinate* fasciculus |
| GCC | *Genu* of *corpus callosum* | TAP (L+R) | Tapetum |

To examine potential group differences in subject motion, we extracted motion parameters from the *ecclog files generated by FSL eddy_correct. These were available for all but the Amsterdam UMC site. Following the approach in Ling et al. (Ling *et al.*, 2012), we examined rotation and translation, each averaged across the X, Y, and Z axes. We found a significant group difference in average rotation (TBI mean=43.9°, control mean=48.2°, $p$=0.018, correcting across motion and translation) and no difference in translation. We conducted follow-up analyses to examine its impact on our results by covarying for rotation. Group differences were consistent when covarying for rotation, indicating that was not a significant confound in our results (**Supplementary Figure 3**). We note that these measures extracted by FSL's tool only quantify motion that is ultimately corrected for, and may not fully capture motion-related artifacts left in the images.

**Supplementary Table 1.** Clinical details and inclusion and exclusion criteria for each of the cohorts included.

| Cohort | Inclusion criteria | Exclusion criteria |
|---|---|---|
| RAPBI + PilotRAPBI | 1) For TBI group: non-penetrating msTBI (moderate-severe) (intake or post-resuscitation Glasgow Coma Scale (GCS) score between 3 and 12); 2) 8-19 years of age; 3) right-handed; 4) normal visual acuity or vision corrected with contact lenses/eyeglasses; and 5) English skills sufficient to understand instructions and be familiar with common words (the neuropsychological tests used in this study presume competence in English). | 1) history of neurological illness, such as prior msTBI, brain tumor or severe seizures; 2) motor deficits that prevent the subject from being examined in an MR scanner (e.g., spasms, movement disorder); 3) history of psychosis, ADHD, Tourette's Disorder, learning disability, mental retardation, autism or substance abuse. These conditions are associated with cognitive impairments that might overlap with those caused by TBI. MRI contraindication was also an exclusion criterion. |
| Baylor (all) | 1) for TBI group: non-penetrating complicated mild or msTBI (moderate-severe) (post-resuscitation GCS score between 3 and 12); 2) 10-18 years of age; 3) right-handed; 4) normal visual acuity or vision corrected with contact lenses/eyeglasses; and 5) fluent in English or Spanish (the neuropsychological tests used in this study were translated into Spanish and administered by bilingual examiners). | 1) history of neurological illness, such as prior msTBI, brain tumor or severe seizures; 2) motor deficits that prevent the subject from being examined; 3) history of psychosis, Tourette's Disorder, learning disability, mental retardation, autism or substance abuse. These conditions are associated with cognitive impairments that might overlap with those caused by TBI. Participants with contraindications to undergoing an MRI scan were excluded. |
| Loma Linda University | 1) age between 4 and 18 years at the time of injury; 2) absence of previous brain injury, neurological disorders, drug or alcohol abuse, or MRI contraindications, including dental braces; 3) moderate-to-severe TBI with Glasgow Coma Scale (GCS) scores between 3 and 12 or complicated mild (cMild) TBI (GCS 13–15) if hemorrhage was detected on an acute brain imaging (CT) exam. Control subjects who met criteria 1 and 2 were recruited from our pediatric clinics and were scanned without sedation. | 1) MRI contraindications; 2) neurological disorders; 3) previous brain trauma or alcohol or drug abuse. |
| Kennedy Krieger | 1) aged 8-18 years of age at time of all study visits; 2) Child and parent are conversant in English, as determined by their ability to understand and participate in English conversations to review the screening form (parent) and consent form (parent and child) 3) For TBI group: moderate or severe traumatic brain injury as defined by first Glasgow Coma Score of 12 or lower upon admission to first emergency room OR post-traumatic amnesia lasting longer than one hour OR alteration in consciousness lasting longer than 15 minutes OR injury-related intracranial abnormality on brain CT or MRI. | 1) children in foster care, 2) penetrating TBI or open TBI (evidenced by dural tear), 3) inability to complete laboratory tasks due to cognitive or motor deficits or uncorrected visual impairment (including red/green colorblindness), 4) ongoing post-traumatic amnesia at the time of first study evaluation or 5) pre-injury diagnosis of mental retardation, psychiatric disorder, or developmental disorder other than ADHD with or without Oppositional Defiant Disorder (ODD). MRI contraindication was also an exclusion criterion. |
| Deakin-1 | All TBI subjects experienced moderate to severe TBI and did no longer participate in acute inpatient rehabilitation. They were tested at least 4 months post-injury when neurological recovery was stabilized. In accordance with the Mayo classification system (Malec et al., 2007), all TBI | Exclusion criteria for both groups were pre-existing developmental disorders, central neurological disorders, intellectual disabilities and musculoskeletal disease. |

| | | |
|---|---|---|
| | patients were classified as 'moderate-to-severe', based on the presence of one or more of the following criteria: loss of consciousness of 30 min or more; worst Glasgow Coma Scale score in the first 24 hours lower than 13, or evidence of contusions, microbleeds or hematoma on CT or MRI images made immediately after the injury. All subjects were able to maintain postural stability during independent stance. | |
| Deakin-2 | The TBI patients were classified as "moderate to severe" based on several factors: the Glasgow Coma Scale score after resuscitation (a subgroup of 8 children had a GCS of 12 or less), the anatomical features of the injury based on inspection by an expert neuroradiologist, and the injury mechanism (traffic accidents and falls), or combinations thereof. All TBI patients were assessed at least 6 months post-injury, when neurological recovery was stabilized. | Participants were excluded if they had pre-existing developmental or intellectual disabilities, a progressive disease, or were taking medication. |
| NCH | 1) hospitalized for at least one night for moderate to severe TBI (post-resuscitation GCS score between 3 and 12) or fracture (control OI group); 2) 8-15 years of age; 3) injured 1-4 years prior. | 1) a history of previous TBI requiring medical treatment (i.e., prior to the target injury); 2) premorbid neurological disorder or mental retardation, or full-time special education placement at school; 3) injury as a result of child abuse or assault; 4) a history of severe psychiatric disorder requiring hospitalization; 5) sensory or motor impairment that precludes completion of study measures; 6) primary language other than English; 7) any contraindication to MRI; 8) refusal by the child's school to participate in the school visit. |
| Amsterdam UMC | Inclusion criteria were: (1) age 8–14 years at time of follow-up; (2) proficient in the Dutch language; (3) children in the TBI group were required to have a history of hospital admission with a clinical diagnosis of moderate/severe TBI (GCS = 12–3, LOC duration >30 min, PTA duration >1 h (Teasdale and Jennett 1976)); and (4) children in the TC group were required to have a history of hospital admission for traumatic injuries below the clavicles (American College of Surgeons 2004). | Exclusion criteria were: (1) previous TBI; (2) visual disorder interfering with neurocognitive testing; or (3) current neurological condition with known effects on neurocognitive functioning, other than TBI, as documented in medical records or reported in a parent-questionnaire for premorbid functioning. |

**Supplementary Table 2.** DTI acquisition parameters for each of the cohorts included.

| Cohort | Scanner | Field strength | Voxel size (mm) | Gradient directions and b-value (s/mm$^2$) | No. b0 volumes |
|---|---|---|---|---|---|
| RAPBI | Siemens TrioTim | 3T | 2x2x2 | 64 at b=1000 | 8 |
| Pilot RAPBI | Siemens Sonata | 1.5T | 2.5x2.5x2.5 | 30 at b=1000 | 5 |
| Baylor-1 | Philips Intera or Achieva | 1.5T | 2.7x2.7x2.7 | 15 at b=860 | 1 |
| Baylor-2 | Philips Intera | 3T | 1.75x1.75x2 | 32 at b=1000 | 1 |
| Baylor-3 | Philips Achieva | 3T | 1.75x1.75x2 | 32 at b=1000 | 1 |
| Loma Linda University | Siemens TrioTim | 3T | 1.2x1.2x3.9 | 60 at b=1000 | 2 |
| Kennedy Krieger | Philips | 3T | 0.83x0.83x2.2 | 32 at b=700 | 1 |
| Deakin-1 | Siemens Magnetom Trio | 3T | 2.2x2.2x2.2 | 64 at b=1000 | 1 |
| Deakin-2 | Philips Intera | 3T | 2x2x2.2 | 45 at b=800 | 1 |
| NCH | Siemens Prisma | 3T | 2x2x2 | 30 at b=700 | 1 |
| UMC Amsterdam | Discovery MR750, GE Healthcare | 3T | 2.5 × 2.5 mm, reconstructed to 1 × 1 × 2.5 mm | 30 at b =750 | 5 |

**Supplementary Video 1.** Significant group differences in the acute/subacute phase. ROIs showing significant group differences are overlaid, with effect sizes corresponding to the color bar.

AcuteDX.mov

**Supplementary Video 2.** Significant group differences in the postacute phase. ROIs showing significant group differences are overlaid, with effect sizes corresponding to the color bar.

PostacuteDX.mov

**Supplementary Video 3.** Significant group differences in the chronic phase. ROIs showing significant group differences are overlaid, with effect sizes corresponding to the color bar.

ChronicDX.mov

**Supplementary Figure 1.** Group differences covarying for rotation parameters. Effect sizes are shown for the primary group comparison in the acute, post-acute, and chronic phases, covarying for sex, age, $age^2$, and average rotation. Cohen's *D* statistics are shown across all midline and bilateral ROIs, along with average FA, with bars indicating the 95% confidence interval. As TBI was coded as "1" and controls as "0", negative effect sizes indicate lower FA in the TBI group. ROI abbreviations are explained in Supplementary Note 1. Dark orange bars indicate significance ($p$<0.005), light orange bars indicate effects that did not withstand multiple comparisons correction (0.05>$p$>0.005), and blue are not significant ($p$>0.05).

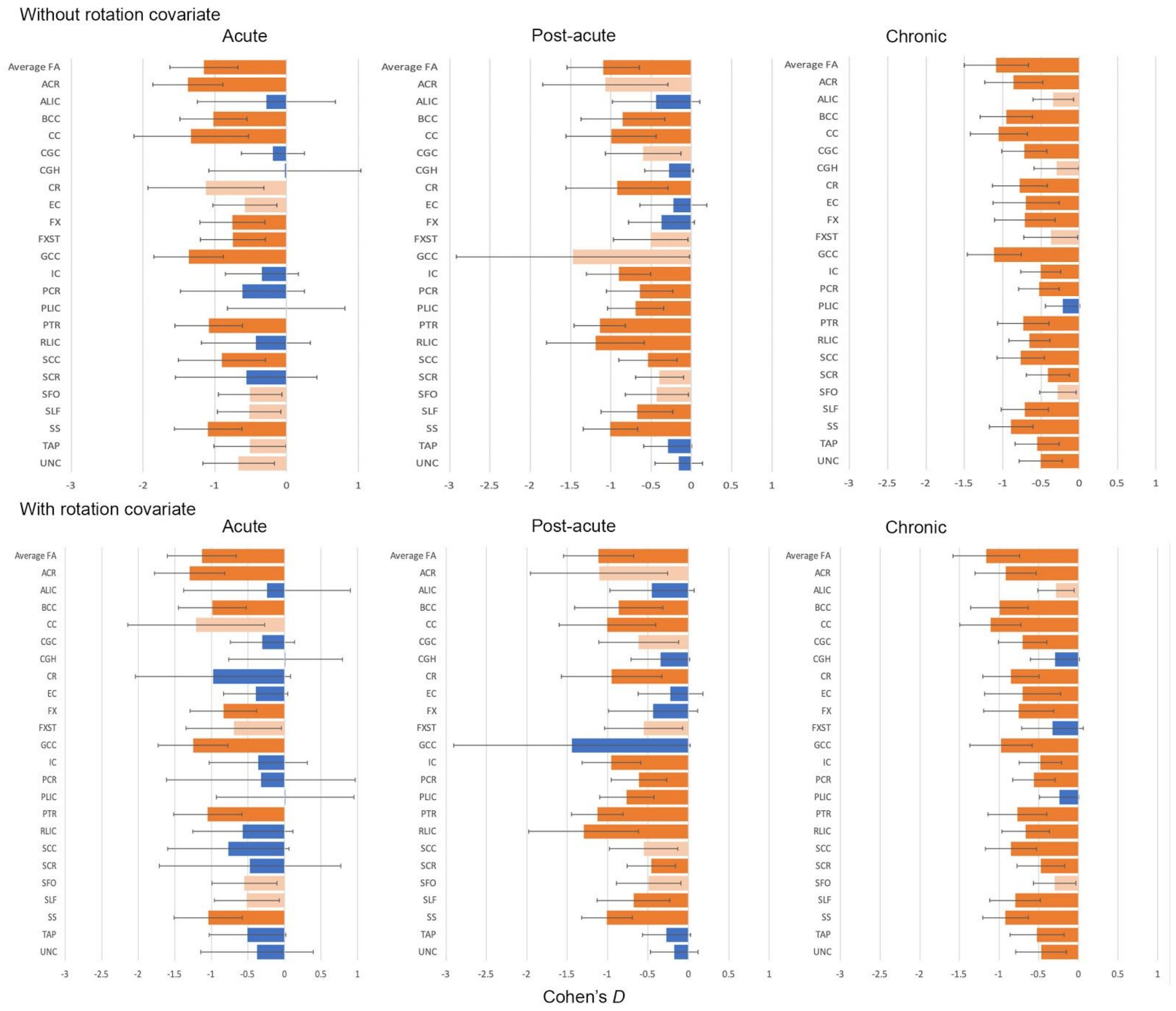

**Supplementary Figure 2.** Group differences in the acute phase for all four diffusion metrics. Effect sizes are shown for FA, MD, RD, and AD, covarying for sex, age, and age$^2$. Cohen's *D* statistics are shown across all midline and bilateral ROIs, along with average metrics, with bars indicating the 95% confidence interval. As TBI was coded as "1" and controls as "0", negative effect sizes indicate lower metrics in the TBI group. ROI abbreviations are explained in Supplementary Note 1. Dark orange bars indicate significance ($p$<0.005), light orange bars indicate effects that did not withstand multiple comparisons correction (0.05>$p$>0.005), and blue are not significant ($p$>0.05).

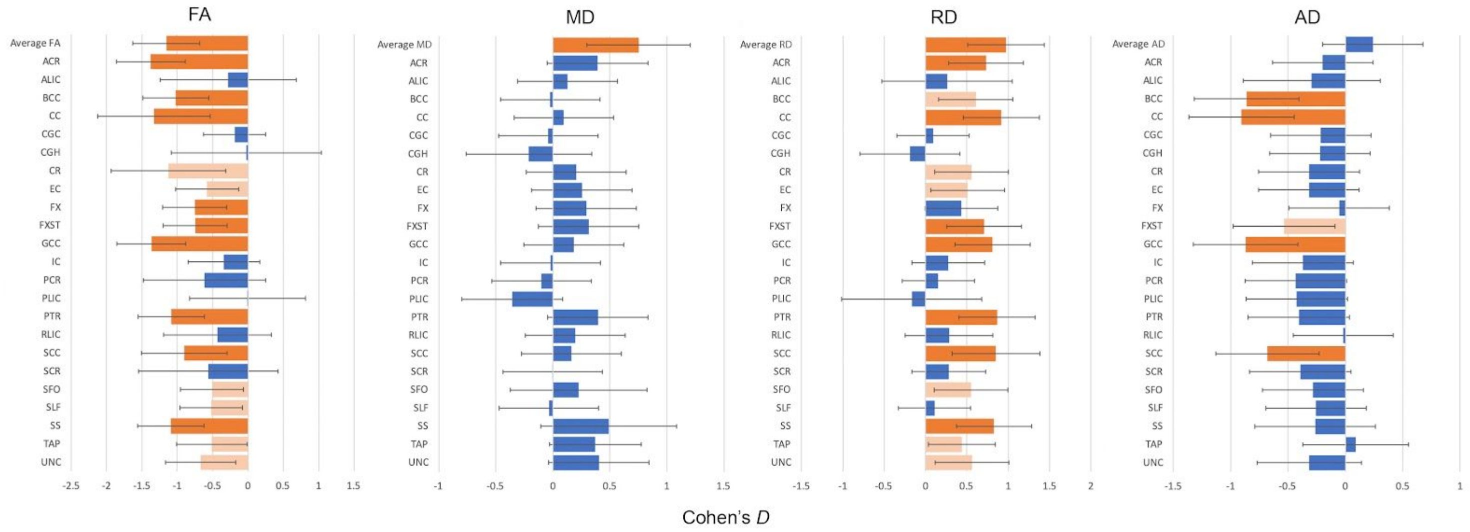

**Supplementary Figure 3.** Group differences in the post-acute phase for all four diffusion metrics. Effect sizes are shown for FA, MD, RD, and AD, covarying for sex, age, and age$^2$. Cohen's *D* statistics are shown across all midline and bilateral ROIs, along with average metrics, with bars indicating the 95% confidence interval. As TBI was coded as "1" and controls as "0", negative effect sizes indicate lower metrics in the TBI group. ROI abbreviations are explained in Supplementary Note 1. Dark orange bars indicate significance ($p$<0.005), light orange bars indicate effects that did not withstand multiple comparisons correction (0.05>$p$>0.005), and blue are not significant ($p$>0.05).

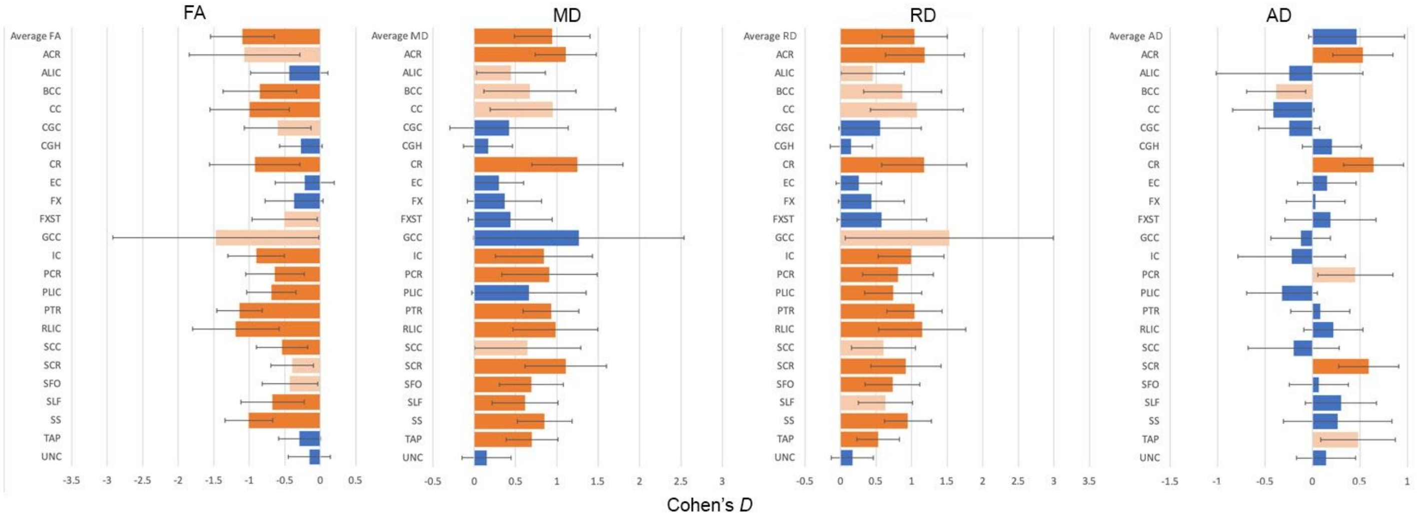

**Supplementary Figure 4.** Group differences in the chronic phase for all four diffusion metrics. Effect sizes are shown for FA, MD, RD, and AD, covarying for sex, age, and age$^2$. Cohen's *D* statistics are shown across all midline and bilateral ROIs, along with average metrics, with bars indicating the 95% confidence interval. As TBI was coded as "1" and controls as "0", negative effect sizes indicate lower metrics in the TBI group. ROI abbreviations are explained in Supplementary Note 1. Dark orange bars indicate significance ($p$<0.005), light orange bars indicate effects that did not withstand multiple comparisons correction (0.05>$p$>0.005), and blue are not significant ($p$>0.05).

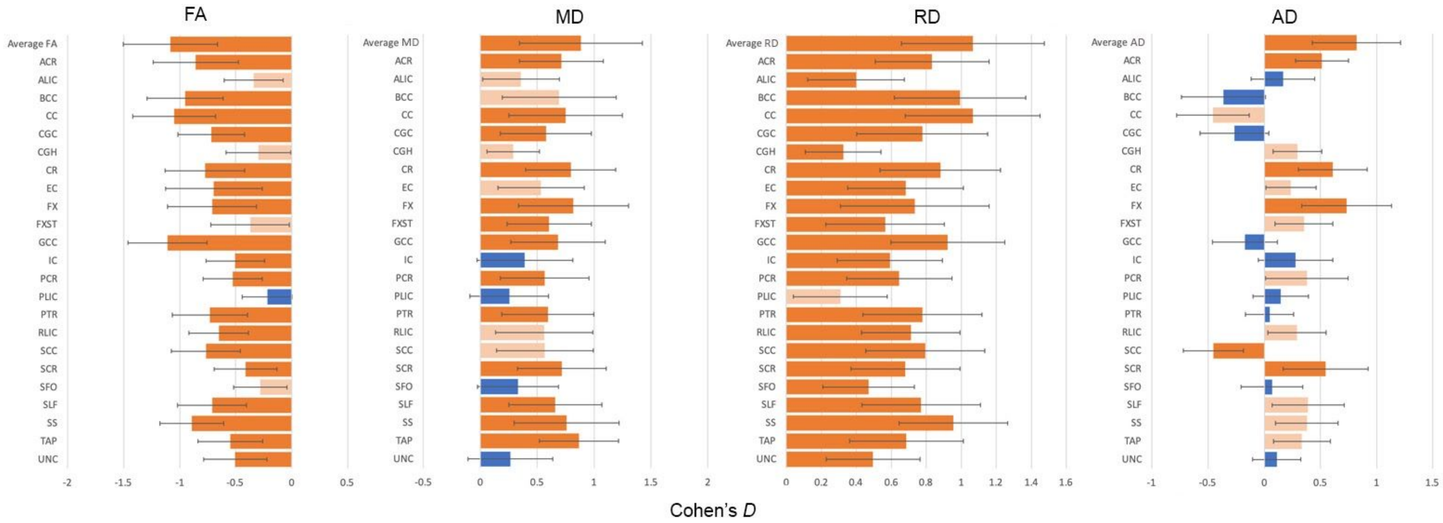

**Supplementary Figure 5.** Group-by-age and group-by-sex interactions for the acute, post-acute, and chronic phases. Unstandardized regression $\boldsymbol{\beta}$s are shown for 23 ROIs and average FA. ROI abbreviations are explained in Supplementary Note 1. Dark orange bars indicate significance ($p<0.005$), light orange bars indicate effects that did not withstand multiple comparisons correction ($0.05>p>0.005$), and blue are not significant ($p>0.05$). Error bars are 95% CI.

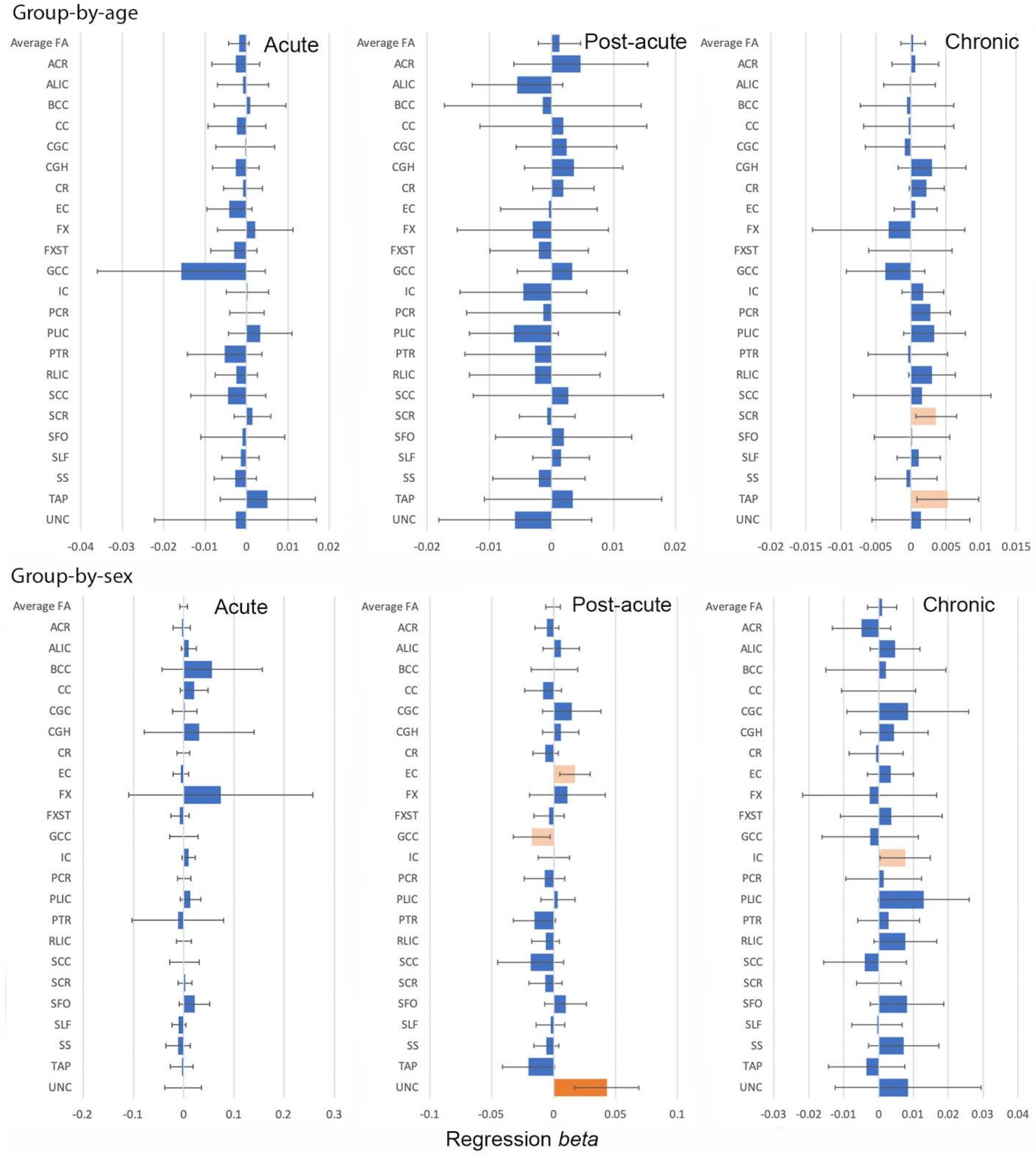

**Supplementary Figure 6.** Group comparisons with healthy controls (HC) and orthopedic injury controls (OI) separated. Effect sizes are shown for FA in the post-acute and chronic phases, covarying for sex, age, and age$^2$. Cohen's *D* statistics are shown across all midline and bilateral ROIs, along with average FA, with bars indicating the 95% confidence interval. As TBI was coded as "1" and controls as "0", negative effect sizes indicate lower FA in the TBI group. ROI abbreviations are explained in Supplementary Note 1. Dark orange bars indicate significance ($p$<0.005), light orange bars indicate effects that did not withstand multiple comparisons correction (0.05>$p$>0.005), and blue are not significant ($p$>0.05).

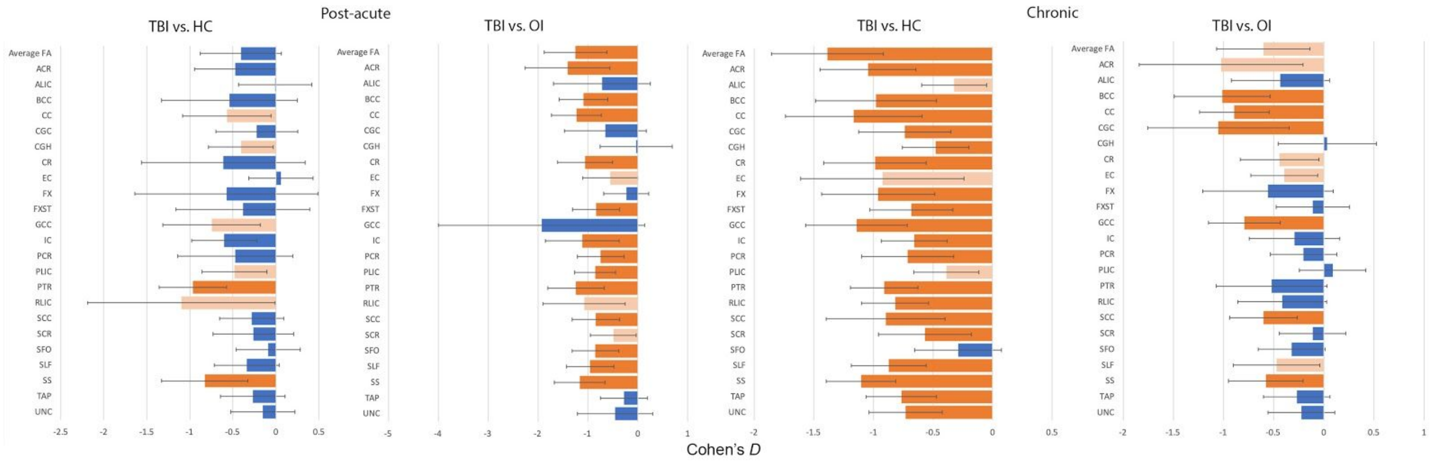

**Supplementary Figure 7.** Linear associations with age-at-injury, time since injury, and Glasgow Coma Scale in the TBI group in the acute, post-acute, and chronic phases. Shown are unstandardized regression ***β***s for 23 ROIs and average FA. ROI abbreviations are explained in Supplementary Note 1. Dark orange bars indicate significance ($p$<0.005), light orange bars indicate effects that did not withstand multiple comparisons correction (0.05>$p$>0.005), and blue are not significant ($p$>0.05). Error bars are 95% CI.

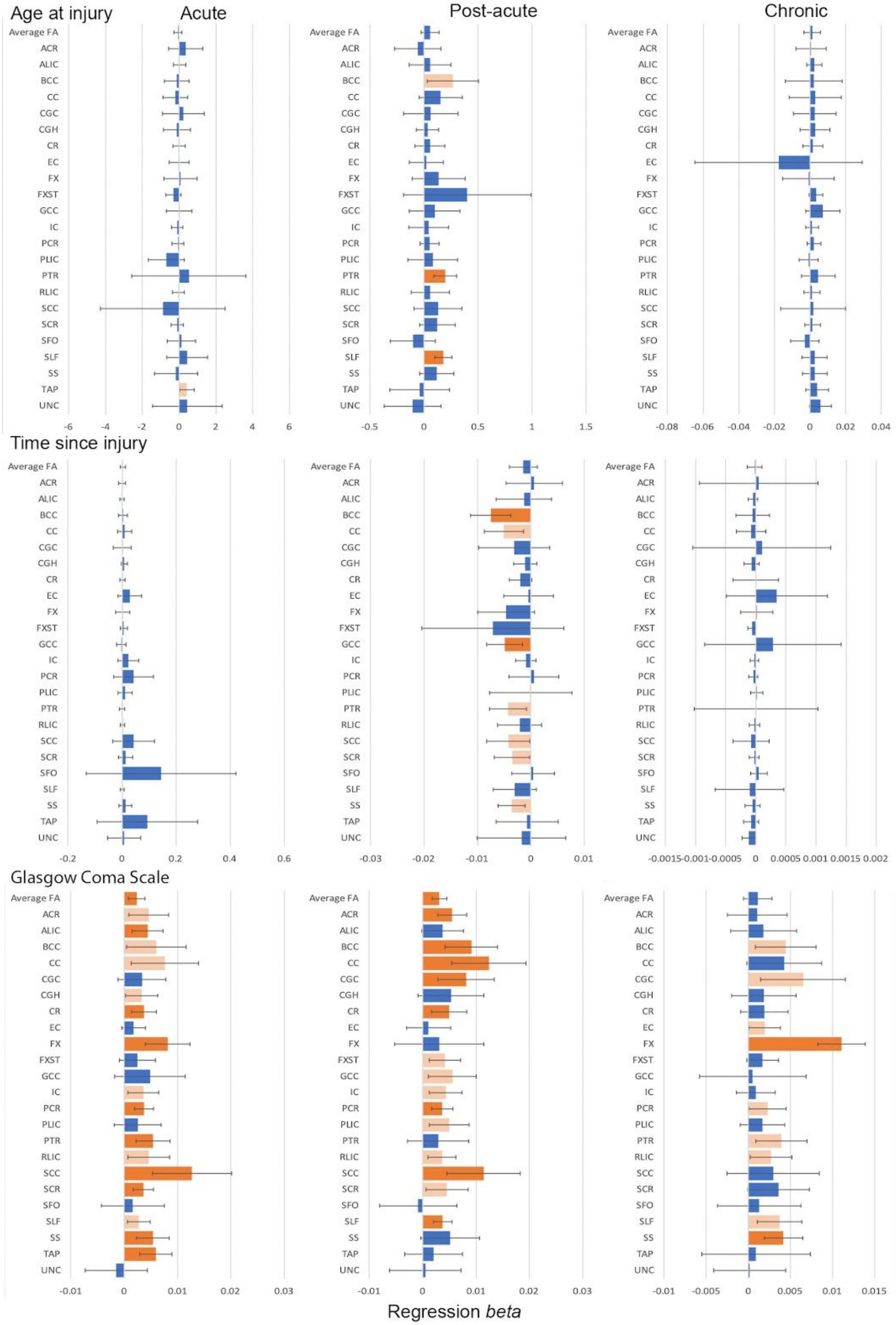

**Supplementary Figure 8.** Linear associations with BRIEF scores in the post-acute and chronic phases. Shown are associations with Global Executive Composite (GEC), Behavioral Regulation Index (BRI), and Meta-Cognition Index (MC). Shown are unstandardized regression $\beta$s for 23 ROIs and average FA. ROI abbreviations are explained in Supplementary Note 1. Dark orange bars indicate significance ($p$<0.005), light orange bars indicate effects that did not withstand multiple comparisons correction (0.05>$p$>0.005), and blue are not significant ($p$>0.05). Error bars are 95% CI.

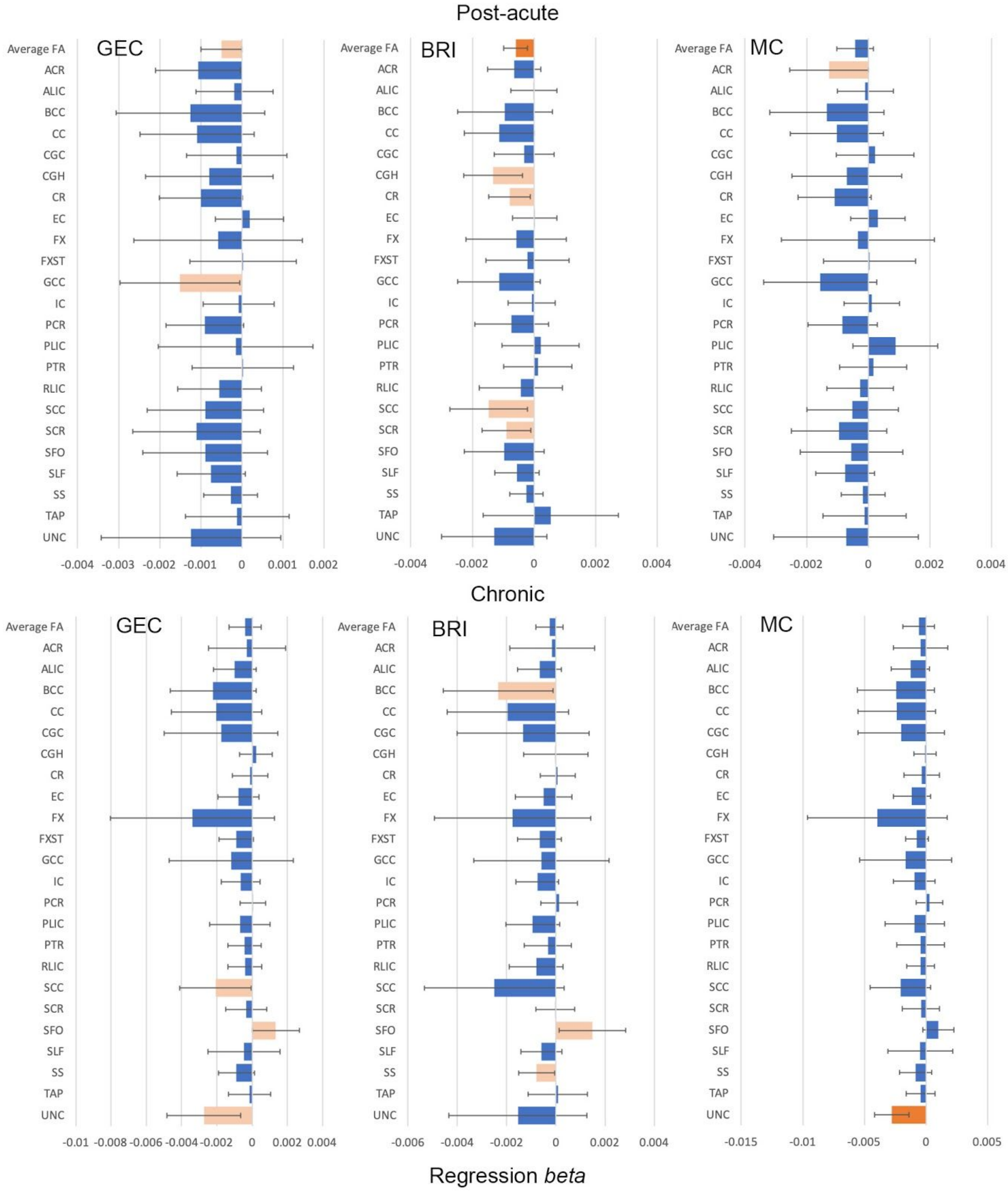
